# Rapidly inducible yeast surface display for antibody evolution with OrthoRep

**DOI:** 10.1101/2024.05.24.595666

**Authors:** Alexandra M. Paulk, Rory L. Williams, Chang C. Liu

## Abstract

We recently developed ‘autonomous hypermutation yeast surface display’ (AHEAD), a technology that enables the rapid generation of potent and specific antibodies in yeast. AHEAD pairs yeast surface display with an error-prone orthogonal DNA replication system (OrthoRep) to continuously and rapidly mutate surface-displayed antibodies, thereby enabling enrichment for stronger binding variants through repeated rounds of cell growth and fluorescence activated cell sorting (FACS). AHEAD currently utilizes a standard galactose induction system to drive the selective display of antibodies on the yeast surface. However, achieving maximal display levels can require up to 48 hours of induction. Here, we report an updated version of the AHEAD platform that utilizes a synthetic β-estradiol induced gene expression system to regulate the surface display of antibodies and find that induction is notably faster in achieving surface display for both our AHEAD system as well as traditional yeast surface display from nuclear plasmids that do not hypermutate. The updated AHEAD platform was fully functional in repeated rounds of evolution to drive the rapid evolution of antibodies.

## Introduction

Orthogonal DNA replication (OrthoRep) is a system for the rapid hypermutation and continuous evolution of chosen genes in *Saccharomyces cerevisiae* [1]. In OrthoRep, an orthogonal DNA polymerase (DNAP) selectively replicates a linear cytoplasmic plasmid (p1) encoding genes of interest at mutation rates up to ∼10^−4^ substitutions per base (s.p.b.) while allowing the genomic error rate to remain at its natural and necessarily low rate of ∼10^−10^ s.p.b. [1, 2]. To enable the rapid evolution of antibodies towards better binding of antigens, we previously coupled OrthoRep with yeast surface display in a system called Autonomous Hypermutation yEast surfAce Display (AHEAD) [3]. In AHEAD, antibody fragments such as nanobodies and scFvs are expressed from p1 as a fusion protein to the yeast agglutinin Aga2. When Aga1 expression is induced from the genome, the rapidly mutating antibody-Aga2 fusion becomes displayed on the yeast surface. The result is a population of yeast autonomously diversifying and displaying antibody variants that can then be guided towards stronger binding through fluorescence activated cell sorting (FACS) (Figure 1a). Successive cycles of culturing and sorting leads to the rapid affinity maturation of antibodies towards desired antigens.

**Figure 1.**
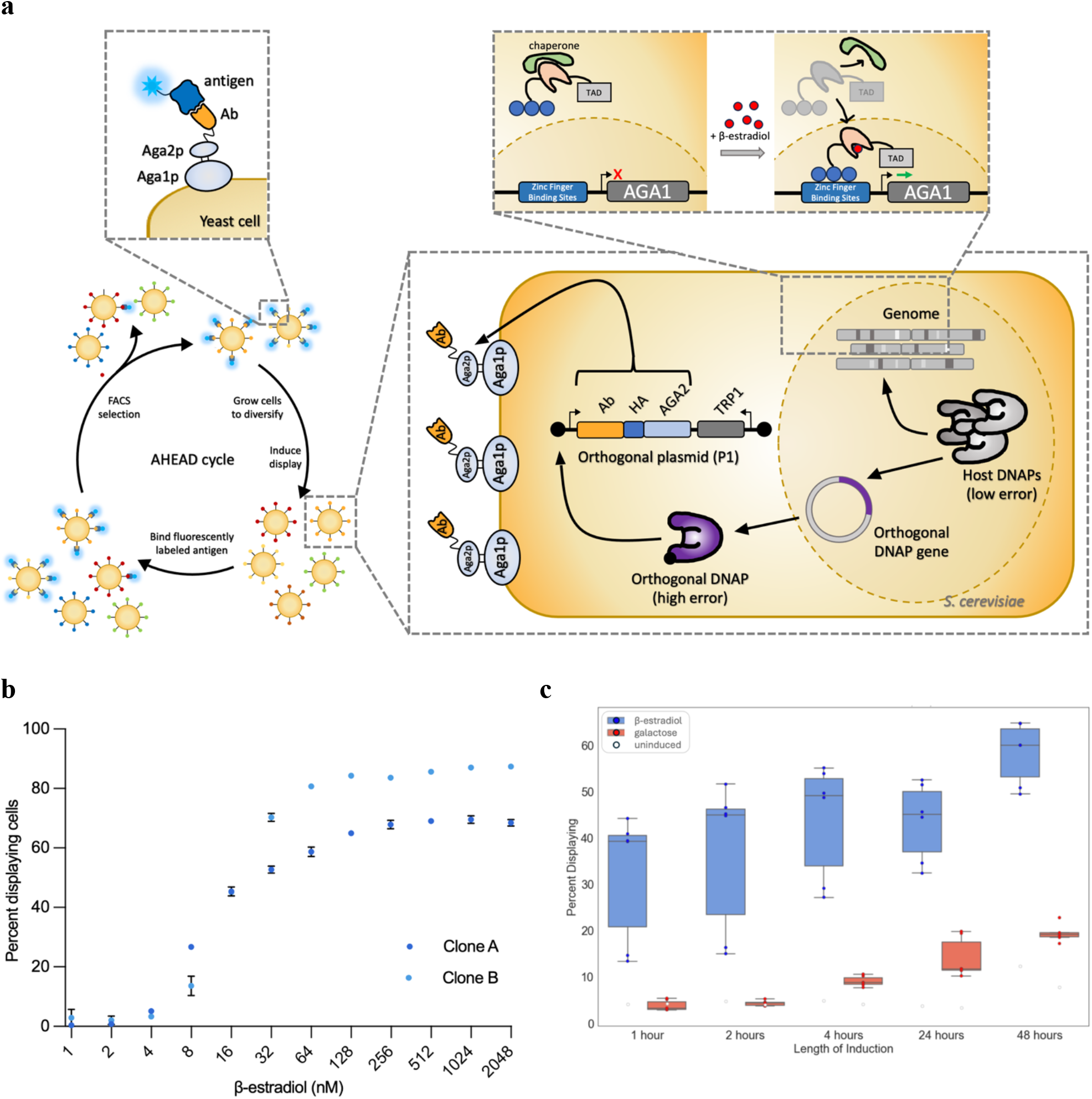
A β-estradiol dependent induction system for AHEAD (**a**) A schematic of an AHEAD cycle for the rapid evolution of binding proteins depicting the mechanism for β-estradiol induction and OrthoRep within yeast cells. (**b**) A dose response curve showing the percent of cells displaying a nanobody after 24 hours of induction at different concentrations of β-estradiol. N=3 for each biological replicate (clone). Points and error bars denote mean and standard deviation. (**c**) The percent of cells displaying a nanobody after 1, 2, 4, 24, and 48 hours of induction in 200 nM β-estradiol induction media. Bars and errors denote maximum, minimum, and quartile ranges. Induction at 30 ºC.

Similar to popular yeast surface display strains such as EBY100 [4], AHEAD relies on galactose induction of Aga1, placed under the control of the *GAL1* promoter *(pGAL1)*. Galactose induction has been extensively studied and is widely used [5-8]. However, this induction method is slow and can require up to 48 hours to achieve maximal surface display levels [9]. Additionally, galactose induction requires a switch into media lacking glucose and is therefore subject to technical idiosyncrasies such as glucose carryover, which acts to represses *pGAL1*. Population-level responses to galactose induction, including time to maximal induction and cell-to-cell variability, are also sensitive to previous growth conditions [10, 11], constituting a “memory” effect that can lead to fluctuation in expression levels achieved with galactose induction.

We reasoned that an alternative induction system could shorten the time required for directed evolution cycles with AHEAD and remove expression idiosyncrasies. During the affinity maturation of antibodies with AHEAD, sorted cells are expanded for 1-3 days depending on the number of cells recovered after the sort [3]. Full induction in galactose can require an additional 2 days, thus making the induction process a ripe target for engineering time-saving alternatives. An ideal induction system would instead allow for rapid induction without extensive manipulation of the media and growth conditions. Human steroid receptors are intracellular, ligand-activated transcription factors that have been successfully reconstituted in yeast to enable the consistent hormone-induced expression of proteins in as little as one hour [12], making them a promising candidate for shortening AHEAD cycles. Moreover, a modular toolkit for the hormone-inducible control of gene expression based on synthetic transcription factors and artificial DNA binding domains responsive to orthogonal hormone inducers was recently reported [13]. In this work, we incorporated the β-estradiol responsive transcription factor and its target promoter (*pER*) from the modular toolkit to drive the expression of Aga1 in AHEAD, resulting in an upgraded AHEAD system. To demonstrate the functionality of the new AHEAD system, we rapidly affinity matured a nanobody against the receptor binding domain (RBD) of SARS-CoV-2’s spike protein.

## Results

### β-estradiol induction of yeast surface display

We constructed an AHEAD strain containing the β-estradiol induction system in which the genomically-encoded Aga1 was placed under the control of *pER* and a nanobody was encoded on p1 as an N-terminal fusion to an HA tag and Aga2 (*i*.*e*., nanobody-HA-Aga2). Upon induction, we observed robust surface display through the detection of the HA tag on cells. We analyzed two important measures of display. First was the proportion of cells displaying nanobodies. In yeast display, it is common to have a bimodal distribution of display where a fraction of cells does not display any detectable protein. Therefore, the proportion of cells displaying nanobodies is arguably the most relevant variable to maximize, as only the displaying cells are subject to selection for functional changes in the displayed nanobody. We found that the percent of cells displaying nanobodies reached its maximum of ∼60%-80% at β-estradiol concentrations exceeding ∼100 nM (Figure 1b). Second was the average level of nanobody display for the displaying subpopulation. It is usually desirable to maximize the level of display within the displaying cells, but arbitrarily high display may be problematic (*e*.*g*., too much display may lead to excessive target depletion when evolving very high affinity binders). We found that β-estradiol concentrations between ∼100 nM and ∼2 mM allowed for a ∼1.5-fold titratable dynamic range in the average level of display within the displaying subpopulation (Figure S1). However, we noticed that concentrations of β-estradiol higher than ∼250 nM negatively impacted cell growth. Thus, we recommend β-estradiol concentrations of ∼100-250 nM for induction in typical cases, as this range achieves the maximum percentage of cells displaying nanobodies with minimal perturbation to cell growth rates. In certain circumstances, it may be desirable to tolerate effects on cell fitness during induction and use higher β-estradiol concentrations to increase the level of display within the displaying population, for example when the sensitivity for detecting target binding of a fluorescently labeled target is low.

### β-estradiol induction speed

Although galactose induction of yeast surface display has been utilized extensively [5-8], maximal surface display can take 48 hours of induction to achieve [9]. We analyzed whether β-estradiol induction can occur faster. When sorting cells during cycles of AHEAD, we aim for a minimum of ∼15-20% of cells displaying the antibody so that sorting bandwidth is not excessively wasted on non-displaying cells. Although both the β-estradiol and galactose inducible strains required 48 hours to reach maximum percentages of the displaying population, the β-estradiol inducible strain approached this maximum much more quickly, reaching sortable levels (∼15-20% of cells displaying) after only 1 hour when induced at 30 °C and 2 hours when induced at room temperature (Figure 1c and Figure S2). This contrasts with the galactose inducible strain, which required 24-48 hours (Figure 1c and Figure S2) to achieve the desired sortable levels. The full induction display levels of the β-estradiol induction system, measured both as percent of cells displaying nanobodies (Figure 1c) and as the average display level of the displaying population (Figure S4a) were comparable or higher than the galactose induction systems in nearly all cases. The sole exception we observed was that the average display level of the displaying population for room temperature galactose induction exceeded that for room temperature β-estradiol induction (Figure S4b). Overall, we conclude that the β-estradiol induction system described here is operationally superior to legacy galactose induction systems for AHEAD, because it achieves full display induction substantially faster than galactose induction without compromising display levels in almost every condition tested. In fact, the percent of cells displaying with the β-estradiol system exceeds that of the galactose induction system at all timepoints (Figure 1c and Figure S2).

The advantage of induction speed of the β-estradiol induction system over galactose induction in the AHEAD system was replicated for nanobody display expressed from a traditional non-hypermutating nuclear CEN/ARS plasmid. However, though we observed substantially faster display induction using β-estradiol and saw a similar percent of cells displaying at maximum induction for both β-estradiol and galactose systems (Figure S3), the β-estradiol induction system achieved a lower average level of display for the displaying subpopulation (Figure S4c and Figure S4d). This discrepancy between the β-estradiol induction system’s performance in the AHEAD system versus a standard nuclear CEN/ARS plasmid display system may be explained by the lower expression levels for the p1-encoded nanobody-Aga2 fusion compared to nuclear expression using *pGAL* or *pER*, which were used to drive the nanobody-Aga2 fusion’s expression in the CEN/ARS plasmid case. In particular, we hypothesize that nanobody-Aga2 expression from p1, but not nanobody-Aga2 expression from the CEN/ARS plasmid, is limiting for both β-estradiol-induced and galactose-induced Aga1 expression. Therefore, the greater strength of galactose-induced Aga1 expression over β-estradiol-induced Aga1 expression can manifest as overall higher display in the CEN/ARS plasmid case – there is enough nanobody-Aga2 to pair with the excess Aga1 – but is throttled by the lower nanobody-Aga2 expression in the p1 case. Nevertheless, the advantages of faster induction times for display with the β-estradiol system will still likely outweigh the lower display levels in standard yeast display from CEN/ARS plasmids for many applications, and researchers should consider using the β-estradiol system presented here even in traditional yeast display systems.

### Nanobody evolution on the updated AHEAD platform

Evolving a binding protein on the AHEAD platform requires repeated cycles of growth, induction, and sorting. To ensure that the updated system utilizing β-estradiol induction could durably withstand repeated cycles of AHEAD, we evolved a low-affinity nanobody “RBD 10” against SARS-CoV-2’s S protein (RBD domain) into a high-affinity nanobody [3]. Evolution of the nanobody was performed with two different error-prone orthogonal DNA polymerases to drive AHEAD’s hypermutation: TP-DNAP1-4-2, the polymerase used in the original publication of AHEAD [3], and “BadBoy3”, a newly developed orthogonal DNA polymerase with an error rate 10-fold higher than TP-DNAP1-4-2 [2]. Throughout AHEAD cycles of evolution, yeast populations behaved as expected, showing robust induction of display and the emergence of new populations representing improved binding to RBD (Figure 2a). After six AHEAD cycles, the final nanobody populations were cloned into a non-hypermutating CEN/ARS plasmid and subjected to two final sorts. Clones were isolated from the high RBD-binding population and sequenced, resulting in the identification of nanobody mutants with 5-6 mutations (Figure 2b and Figure S5). These evolved variants were characterized by on-yeast EC_50_ measurements and found to have an EC_50_ of 10.3nM and 3.2 nM, in comparison to the parent clone’s *K*_D_ of 417 nM (Figure 2c) [3]. Indeed, AHEAD with the new β-estradiol induction is functional and capable of generating high-affinity nanobodies through a rapid evolutionary process.

**Figure 2.**
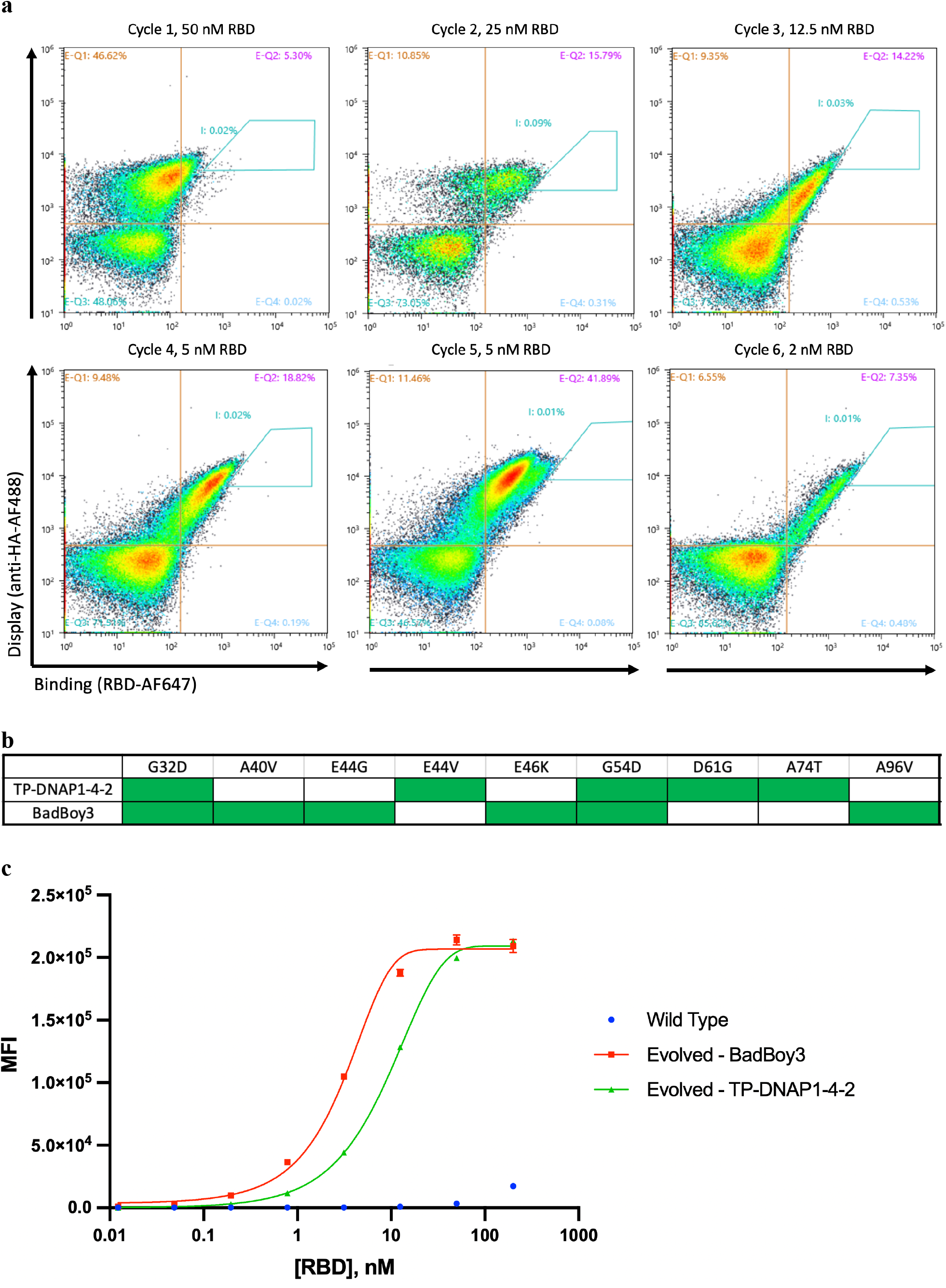
Successful evolution of nanobodies in a β-estradiol inducible AHEAD strain. (**a**) FACS plots for six cycles of AHEAD using the error-prone DNA polymerase TP-DNAP1-4-2 for hypermutation. Blue gates indicate cells sorted to seed the next cycle. (**b**) Mutations found in top variants isolated from both nanobody evolution campaigns carried out with two different OrthoRep DNA polymerases. Naming refers to the DNA polymerase used for the evolution. (**c**) On-yeast EC_50_ curves showing increase in apparent binding affinity of evolved clones compared to parent (wild type). Evolved clones were encoded on a standard non-hypermutating CEN/ARS plasmid for on-yeast EC_50_ curves. N=3 for each measurement. Points and error bars denote standard deviation. For some points, the error bars are obscured because they are too close to the mean.

## Discussion and Conclusion

We have engineered a new version of AHEAD that relies on a synthetic β-estradiol-inducible expression control system [13] to drive yeast surface display of nanobodies. This improvement shortens the time required for induction of surface display from up to 48 hours to as little as 1 hour, enabling a FACS sort to be completed on the same day as induction. One cycle of AHEAD can now be reliably performed within 2-3 days – or shorter if more cells are recovered when sorting – comparable to the time required only for cells to reach saturation following sorting in the initial version of AHEAD. We further showed that the new induction system’s performance is robust over repetitive cycles in a full AHEAD affinity-maturation campaign. In addition to time savings for AHEAD, the new induction system has greater ease of use than galactose induction, as it requires no preparation of separate growth media or washing steps prior to induction. The new induction system has advantages over traditional galactose induction systems for standard display platforms with CEN/ARS plasmids as well.

## Associated Content

### Supporting Information

#### Materials and Methods

Figure S1: Median fluorescence intensity of displaying cells at varying β-estradiol concentrations

Figure S2: Induction time course at room temperature, nanobody on p1

Figure S3: Induction time courses in CEN/ARS format

Figure S4: Median fluorescence intensity of displaying cells

Figure S5: Sequencing of post-evolution individual clones

Table S1: Key strains used in this study

Table S2: Key plasmids used in this study

## Abbreviations

AHEAD: autonomous hypermutation yeast surface display
FACS: fluorescence activated cell sorting
DNAP: DNA polymerase
RBD: receptor binding domain

## Author Contributions

A.M.P., R.L.W., and C.C.L. designed the experiments. A.M.P. conducted the experiments. A.M.P. and C.C.L. analyzed the data and wrote the manuscript.

## Acknowledgements

This work was funded by NIH R01CA260415 and NIH R35GM136297 to C.C.L. Training grant support to A.M.P. was provided by NIH T32GM136624. R.L.W. is supported by a Hewitt Foundation for Medical Research Postdoctoral Fellowship. We thank the Institute for Rapid Antibody Engineering and Evolution, part of the Engineering+Health Initiative of the UCI Samueli School of Engineering, for additional support.

## References

(1) Ravikumar, Arjun et al. (2018) Scalable, Continuous Evolution of Genes at Mutation Rates above Genomic Error Thresholds. Cell vol. 175,7

(2) Rix, Gordon et al. (2023) Continuous evolution of user-defined genes at 1-million-times the genomic mutation rate. bioRxiv. Preprint.

(3) Wellner, Alon et al. (2021) Rapid generation of potent antibodies by autonomous hypermutation in yeast. Nature chemical biology vol. 17,10

(4) Boder, E T, and K D Wittrup. (1997) Yeast surface display for screening combinatorial polypeptide libraries. Nature biotechnology vol. 15,6

(5) Lohr, D et al. (1995) Transcriptional regulation in the yeast GAL gene family: a complex genetic network. FASEB vol. 9,9

(6) Barnett, James A. (2004) A history of research on yeasts 7: enzymic adaptation and regulation. Yeast vol. 21,9

(7) Sellick, Christopher A et al. (2008) Galactose metabolism in yeast-structure and regulation of the leloir pathway enzymes and the genes encoding them. International review of cell and molecular biology vol. 269

(8) Traven, Ana et al. (2006) Yeast Gal4: a transcriptional paradigm revisited. EMBO reports vol. 7,5

(9) Zhao, Jing-Zhuang et al. (2017) An efficient and simple method to increase the level of displayed protein on the yeast cell surface. Journal of microbiological methods vol. 135

(10) Zacharioudakis, Ioannis et al. (2007) A yeast catabolic enzyme controls transcriptional memory.” Current biology: CB vol. 17,23

(11) Stockwell, Sarah R et al. (2015) The yeast galactose network as a quantitative model for cellular memory. Molecular bioSystems vol. 11,1

(12) Poletti, A et al. (1992) A novel, highly regulated, rapidly inducible system for the expression of chicken progesterone receptor, cPRA, in Saccharomyces cerevisiae. Gene vol. 114,1

(13) Sanford, Adam et al. (2022) A Toolkit for Precise, Multigene Control in Saccharomyces cerevisiae. ACS synthetic biology vol. 11,12

(14) Gunge, N. & Sakaguchi, K. (1981) Intergeneric transfer of deoxyribonucleic acid killer plasmids, pGKL1 and pGKL2, from Kluyveromyces lactis into Saccharomyces cerevisiae by cell fusion. J. Bacteriol. 147, 155–160

