## Supplementary Information for "Rapidly inducible yeast surface display for antibody evolution with OrthoRep"

### Methods

#### Strain and plasmid construction

For a list of key strains and plasmids, see Table S1 and Table S2. For galactose induction, previously published strains EBY100 (ATCC MYA-4941) and yAW301 [3] were used. β-estradiol inducible versions of these two strains were constructed by genomically integrating *AGA1* under the control of *pER* [13] at the *AGA1* locus. The synthetic transcription factor that responds to β-estradiol (synER [13]) to induce expression was genomically integrated at the *URA3* locus, resulting in yeast strain yAP174. A protoplast fusion [14] using an F102-2 strain harboring a "landing pad" p1 [3] as the donor strain was performed to incorporate the p1 and p2 plasmids into yAP174. The resulting strain was transformed with the error-prone polymerase TP-DNAP1-4-2 or BadBoy3 encoded on a CEN/ARS plasmid (pAW038 and pAW729), yielding strains yAP193 and yAP196, respectively. In the galactose strains, *AGA1* is expressed under the control of *pGAL* at the *AGA1* locus and the synthetic transcription factor is not present, but strains were identical in all other regards. To test induction of yeast surface display, a nanobody (RBD10) [3] fused to an HA tag was either expressed from a CEN/ARS plasmid under *pGAL* or *pER* in EBY100 (galactose induction) and yAP174 (β-estradiol induction) or from p1 in yAW301 (galactose induction) and yAP193/yAP196 (β-estradiol induction).

Nanobody RBD10 was cloned into pAP175, pAP176, and pAP191 using Golden Gate Assembly. pAP175 and pAP176 are CEN/ARS plasmids for yeast surface display driven by *pGAL* and *pER*, respectively. pAP191 is a plasmid into which genes of interest can be encoded as fusions to AGA2 with *TRP1* and NatMX selection markers. Linear DNA for with homologous flanks for integration onto the p1 landing pad is prepared by digestion of pAP191 derived plasmids with ScaI, or alternatively by PCR amplification.

#### β-estradiol induction

Nanobody RBD10 was integrated onto the p1 landing pad [3] in yAP193 using a ScaI-linearized pAP191 containing the nanobody, and *TRP1* and NatMX were used to select for the recombinant p1. Two colonies were picked into liquid media selecting for *TRP1* and grown to saturation. 24 well blocks were prepared to contain serial dilutions of β-estradiol in SC-HLUW (Synthetic Complete media lacking Histidine, Leucine, Uracil and Tryptophan) and saturated yeast cultures were passaged 1:100 into the induction media blocks such that each of the two cultures would be induced in triplicate. Passaged cultures were grown for 24 hours with shaking (200rpm) at 30 ºC. Approximately $10^6$ cells from each culture were transferred to a 96 well plate, washed with sorting buffer (20 mM Tris-HCl pH 7.5, 100 mM NaCl, 0.1% BSA and 5 mM maltose), and stained for 30 minutes at 4 ºC with shaking in 100 uL primary staining solution containing buffer with 13.3 nM mouse-anti-HA antibody (gift from A. Kruse). Cells were washed twice in sorting buffer and stained for 15 minutes at 4 ºC with shaking in 100 uL secondary staining solution containing buffer with 13.3 nM goat-anti-mouse-PE antibody (Abcam ab97024). Cells were washed twice more and resuspended in 250 uL sorting buffer and display signal was measured on an Attune NxT cytometer (Thermo Fisher).

We found that levels of display induced could vary across clones when expressed from p1, which we have previously observed and attribute to early differences in p1 copy number after the integration of the nanobody expression construct onto p1, since it takes time for p1 copy number to reach steady state.

**Induction time course measurements**

Nanobody RBD10 was transformed into EBY100 and yAP174 in CEN/ARS format (in pAP175 and pAP176, respectively) or integrated onto the p1 landing pad in yAW301 [3] and yAP193 using pAP191. *TRP1* was used to select for successful transformants in the CEN/ARS format and *TRP1* and NatMX were used to select for successful transformants in the p1 format. 2-3 colonies from each transformation were grown in liquid media selecting for *TRP1*. When cultures reached an $OD_{600}$ between 2.0-3.0, they were passaged 1:20 into corresponding induction media as well as a non-induced control. For each strain, each of the 2-3 biological replicate cultures was induced in 2-3 replicates. For β-estradiol inductions, a concentration of 200 nM β-estradiol was used. Cells were grown at 30 ºC and $10^6$ cells were removed for display measurements after 1, 2, 4, 24, and 48 hours. The experiment was also repeated with growth at room temperature (Figure S2, Figure S3b and Figure S4 b,d). Cells were stained and measured as described in the previous section.

**Evolution of anti-RBD nanobodies**

A colony from the transformation and integration of RBD10 onto p1 using yAP193 was picked into liquid SC-HLUW and grown to saturation. The culture was passaged 1:100 into SC-HLUW with 200 nM β-estradiol and grown overnight (18-24 hours) at 30 ºC with shaking at 200 rpm. Approximately $5x10^7$ cells were taken from the induced culture, washed with sorting buffer, and stained for 1 hour at 4 ºC with rotation in 250 uL of a primary staining solution containing buffer with 50 nM RBD-AF647 (Acro Biosystems. Labeled with AF647 NHS Ester (Thermo Fisher) in-house) and 20 nM anti-HA-AF488 (R&D systems). Cells were washed twice with sorting buffer and resuspended in 4 mL buffer. Cells were sorted into SC-HLUW using FACS (SONY SH800), selecting 250 cells from the gate (gates shown in Figure 2a). Note that cell size changes upon exposure to β-estradiol, so attention should be paid to FSC v SSC sorting gates. Sorted cells were grown at 30 ºC with shaking (200 rpm) until saturation, and the process of induction, staining, sorting, and growth was repeated for a total of 6 rounds. RBD concentration was reduced each round except for round 5 (Figure 2a). The evolution was repeated in a second strain, yAP196, containing a newer error-prone DNA polymerase, "BadBoy3" [2]. The p1 sequences from the final cycle's selected population were amplified by PCR following a yeast miniprep and cloned into a CEN/ARS display plasmid library. This library was transformed into yAP174 and the subsequent populations were sorted twice more on FACS to isolate the best binding clones. This sorted population was plated and ten colonies at random were sequenced (Figure S5). The most highly represented sequence from each evolution experiment was chosen for further characterization.

**On-yeast $EC_{50}$ curves**

Representation among the ten colonies sequenced at the end of each evolution experiment was taken as a proxy for binding strength, and a clone of the most represented sequence from each experiment was chosen to measure binding strength via on-yeast $EC_{50}$ curves. Separate cultures of yeast containing plasmids for the β-estradiol-inducible expression of each selected evolved nanobody, as well as the wild type nanobody, were grown to $OD_{600}$ = 2.0-3.0 in selective media and induced by 1:100 passaging into SC-HLU with 200 nM β-estradiol and grown overnight at 30 ºC with shaking. Cultures were pelleted by centrifugation at 4 ºC, 750 rpm, washed with sorting (HBSBM) buffer (20 mM Tris-HCl pH 7.5, 100 mM NaCl, 0.1% BSA and 5 mM maltose), and resuspended in 1 mL HBSBM. Samples of the cultures were diluted and measured on a flow cytometer to obtain precise cell densities of the cell suspensions, and 5 mL of cells in HBSBM at a density of $10^6$ cells per mL were prepared. Serial dilutions of biotinylated antigen (SARS-CoV-2 spike RBD, Acro Biosystems, biotinylated by the NHS ester method in-house) were prepared and 100 uL of each dilution was dispensed into wells of a 96-well block such that each antigen concentration for each nanobody clone selected would be measured in triplicate. To this, 100 uL of cell suspension of each clone was added. The plate was incubated with shaking (200 rpm) at 4 ºC for one hour, washed with HBSBM, and pellets resuspended in 50 uL secondary binding solution containing 1 uL anti-HA-AF488 antibody (R&D Systems) and 1 uL streptavidin-PE (BD Biosciences) per mL of solution, in HBSBM. After 15 minutes of secondary binding solution incubation with shaking at 4 ºC, cells were washed again with HBSBM and resuspended in 150 uL HBSBM buffer. Display and binding were measured and recorded on a flow cytometer. Displaying populations were gated and the mean fluorescence intensity of binding within the displaying populations was plotted against the antigen concentration.

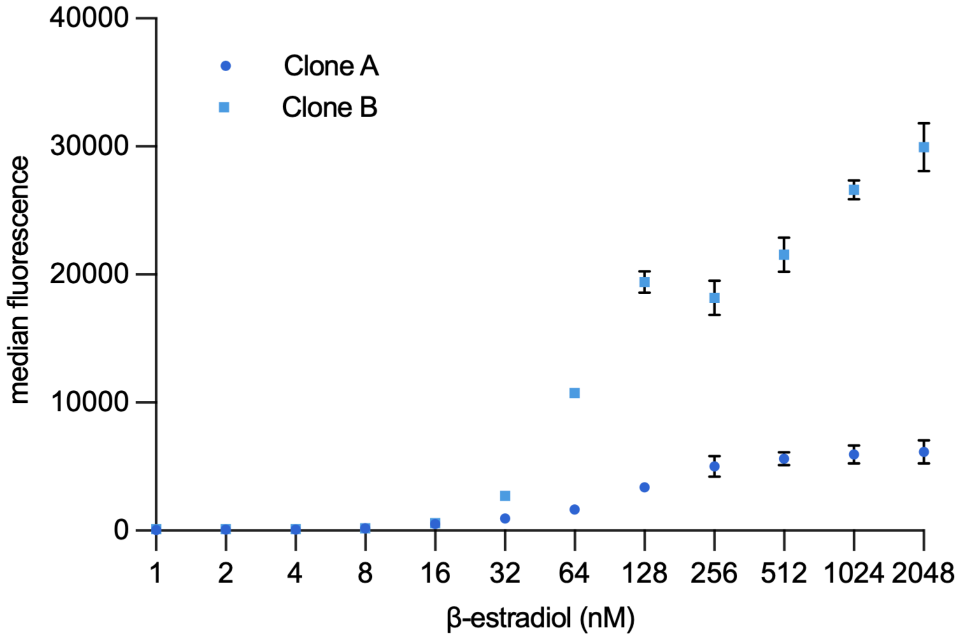


**Figure S1:** Median fluorescence intensity of displaying cells captured under "percent displaying cells" in Figure 1b. At concentrations resulting in maximum percent of cells displaying (>100 nM), there is a titratable range where per-cell display can be increased ~1.5 fold. N=3 for each biological replicate (clone); points and error bars denote mean and standard deviation. We note the difference in display between the two clones, which we attribute to differences in the copy number of p1-encoded nanobody at time of measurement. The copy number difference can occur early after integration of the nanobody expression construct onto p1, as a single integrant increases its copy number within cells over time to reach a steady-state concentration.

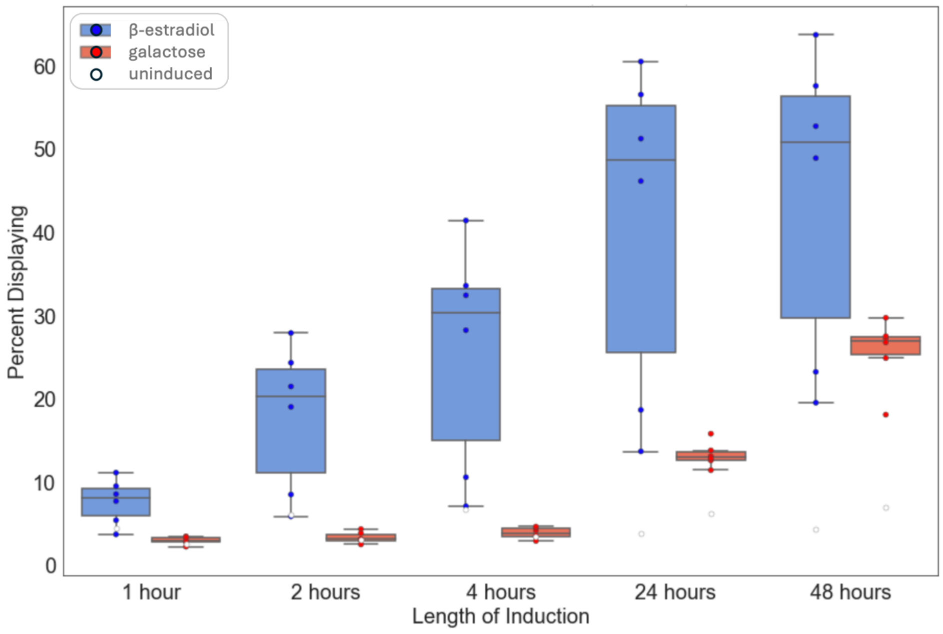


**Figure S2.** Induction of surface display of a nanobody on AHEAD at room temperature. The percent of cells displaying a nanobody after 1, 2, 4, 24, and 48 hours of induction in 200 nM β-estradiol induction media. Induction at room temperature (~22 ºC). For induced samples, N=6, composed of 3 biological replicates (clones) with 2 technical replicates each. Bars and errors denote maximum, minimum, and quartile ranges.

a

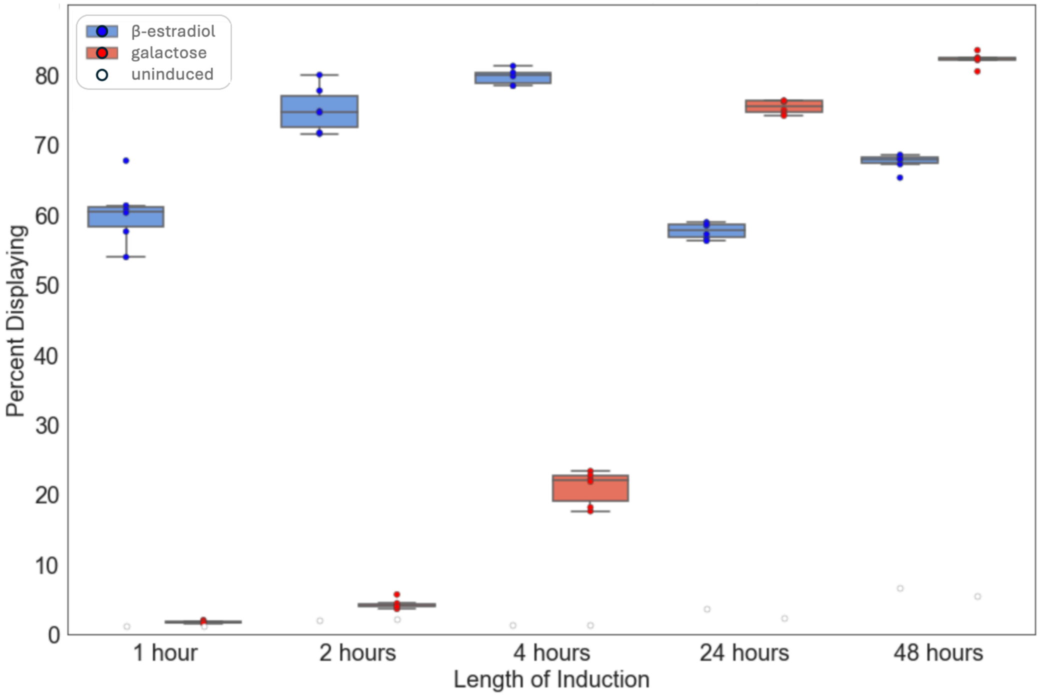


b

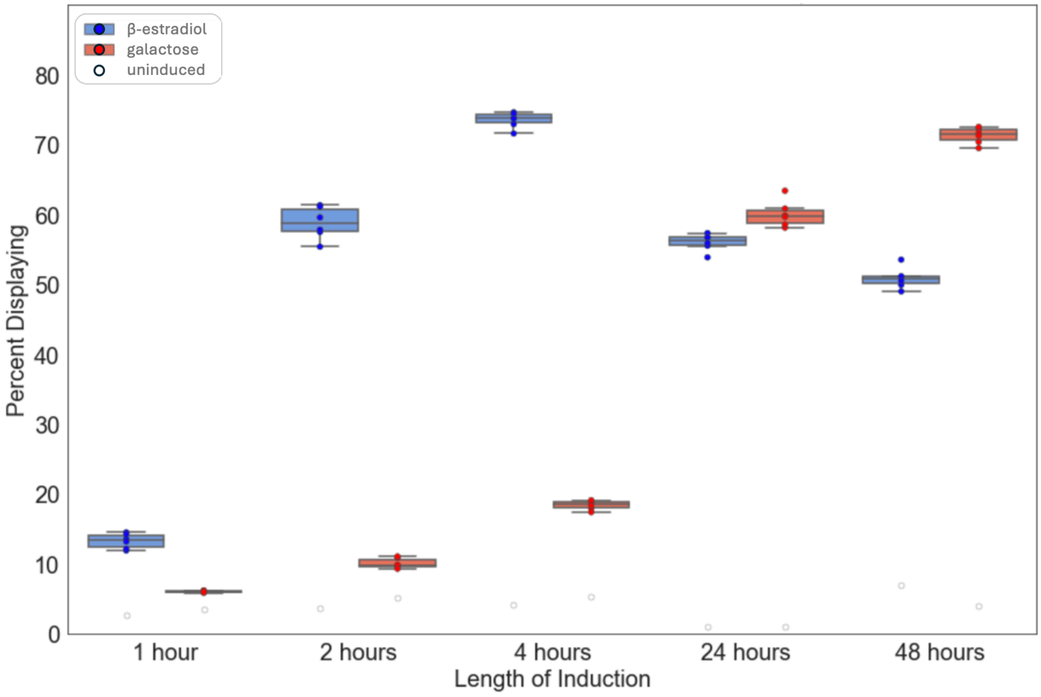


**Figure S3.** Induction of surface display of a nanobody on a CEN/ARS plasmid. Percent of cells displaying a nanobody after 1, 2, 4, 24, and 48 hours of induction in 200 nM β-estradiol induction media. For induced samples, N=6, composed of 3 biological replicates (clones) with 2 technical replicates each. Bars and errors denote maximum, minimum, and quartile ranges. (**a**) Induction at 30 ºC. (**b**) Induction at room temperature (~22 ºC).

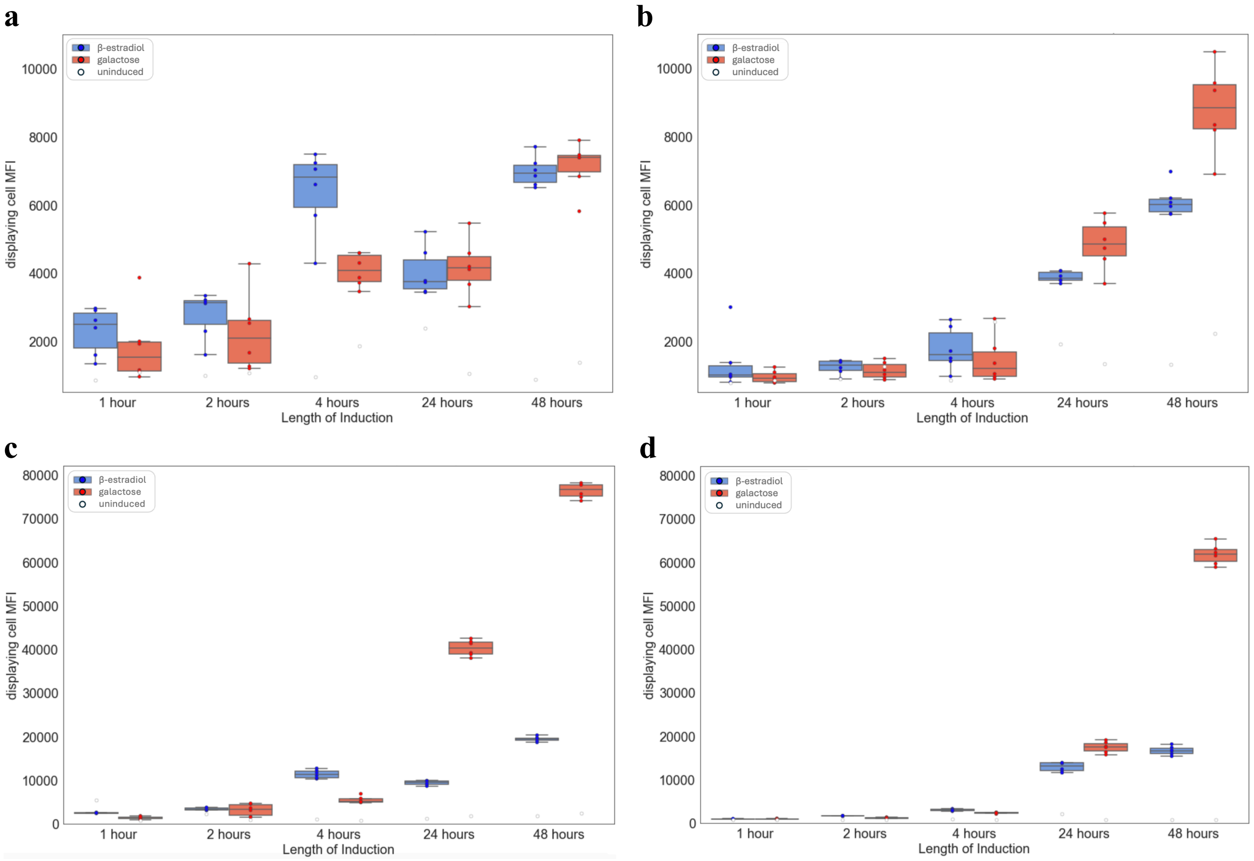


**Figure S4:** Median fluorescence intensity of displaying population. For induced samples, N=6, composed of 3 biological replicates (clones) with 2 technical replicates each. Bars and errors denote maximum, minimum, and quartile ranges. **(a)** Nanobody expressed from p1, 30 ºC induction. **(b)** Nanobody expressed from p1, room temperature (~22 ºC) induction. **(c)** Nanobody expressed from CEN/ARS, 30 ºC induction. **(d)** Nanobody expressed from CEN/ARS, room temperature (~22 ºC) induction.

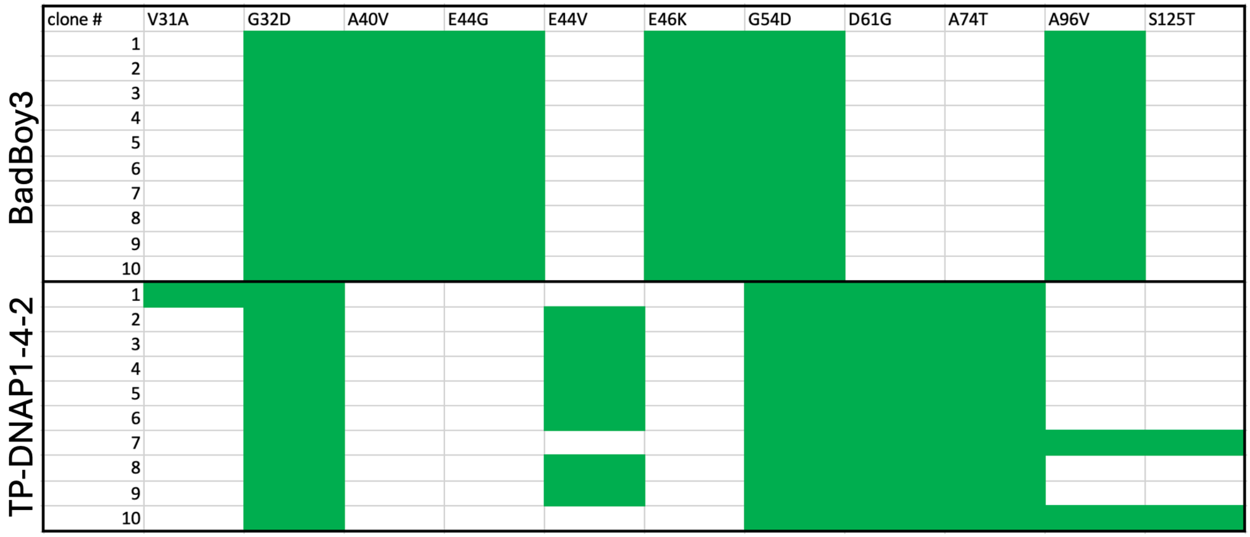


**Figure S5.** Sequencing of ten individual post-evolution colonies. Green boxes indicate presence of the mutation. Polymerases used during the evolution are listed on the left. Clone #3 of both evolutions was chosen for on-yeast $EC_{50}$ characterization.

**Table S1.** Key strains used in this study

| Yeast | Description | Genome |
|---|---|---|
| yAW301 | galactose inducible AHEAD strain with TP-DNAP1-4-2 polymerase encoded on a CEN/ARS nuclear plasmid | MATa AGA1::pGAL1-AGA1-URA3 ura3-52 trp1Δ leu2-delta200 his3delta200 pep4::HIS3 prb1delta1.6R can1 GAL MET15::KanMX; p1-MET15 landing pad; pAW038 |
| yAP193 | β-estradiol inducible AHEAD strain with TP-DNAP1-4-2 polymerase encoded on a CEN/ARS nuclear plasmid | MATa AGA1::pER-AGA1-HygR ura3-52::synTF-URA3 trp1Δ leu2delta1 his3delta200 pep4::HIS3 prb1delta1.6R can1 GAL met15Δ; p1-MET15 landing pad; pAW038 |
| yAP196 | β-estradiol inducible AHEAD strain with BadBoy3 polymerase encoded on a CEN/ARS nuclear plasmid | MATa AGA1:: pER-AGA1-HygR ura3-52::synTF-URA3 trp1Δ leu2delta1 his3delta200 pep4::HIS3 prb1delta1.6R can1 GAL met15Δ; p1-MET15 landing pad; pAW729 |
| EBY100 | galactose inducible yeast surface display strain | MATa AGA1::pGAL1-AGA1-URA3 ura3-52 trp1Δ leu2delta200 his3delta200 pep4::HIS3 prb1delta1.6R can1 GAL |
| yAP174 | β-estradiol inducible yeast surface display strain | MATa AGA1::pER-AGA1-HygR ura3-52::synTF-URA3 trp1Δ leu2delta1 his3delta200 pep4::HIS3 prb1delta1.6R can1 GAL |

**Table S2.** Key plasmids used in this study.

| Plasmid name | Description | Sequence |
|---|---|---|
| pAW038 | CEN/ARS plasmid encoding TP-DNAP1-4-2 DNA polymerase with a LEU2 marker for yeast and a KanR marker for propagation in *E. coli*. | gagctcgtgttgttatccgatacaaccggatatttttcttttaatgagtctaaaccgtgatagcttcaggttaatacaatcaaaaaaagctcaaatattcttttaatgccgcgttcacagattccaattgaatacaactaggtagttcattatatgaagcctttgctactatttttcactatagtctgccttcaccttaatgcagacatccacatattttaatcactttaaaataaaaaggaagatatattagaagctatgatccaatctgtaagccagattaaaattcacgaactcttctttcatttgaattgaatgctttgagttggggtagattatcgcaaattactcatcacatttattgactacgaacttgctgatgtcctttttttatttatatttttcttcagtgaagcgatttttttttacacagaccaagacggaaaaaagtagctaaggaagaaaacaaaatcatgaaaaaaatgtgaagtgatcatgcacatcgcatcaacttaaacattggcttagagatatatagagttagagtttacggcaacctttaagcaccaatacctttttggcatagtctaaagacctggttcttaattttaaacaaatttaactaaagatttccctatcaaagaagtaacgagttgacagattttctcaaaataaatcgatactgcatttctaggcatatccagcgagatctatggactacaaagacaaagcattaaacgacttgagaaacgtatacgcagacttcgactccttgccattggacttcagacaaatcttaattaaggatagagctacattgttgcaaaaggaagacgttgaaaagaaaattttggaaagacaagaagatgccaaaaagtacgctgaatacttgaagcaatcagaaatcccagaaagaatctccttacctaacatcaagagacataagggtgtatcaatctcctttgaagaaacatctgaagatatggtcttggaaccaagaccttttatcttcgacggtttgaacatcagatgtttcagaagagaaaccatattcagtttgaaaaataagattttgaacatggttaaggaatcttcatcctttaaaaacgtttcaagacaaagtgtatcttttatgtacttcaaaatttttaataagggtaaagttattgcaagtacaaagtctgttaacatctatgaagataagatcgacgaaagattggaagatttgtgtaacaacttcgatgacgtattgaaaaagattatagacgtcacctatggttacgaatccttattcgttagtgaaacttactcttacgtaatattctatgctaaatcaatctactttccacaacctagatgcgttaacaactggggtaacaacatcccaaacatcttgacattcgattcctttaagcttttcaccgcaaacaaaaacaacgttagttgtatcaagcaatgttcaagatttttgtggcaaaaagatttcaacactttagaagaaatgatagaatacaagaacggtaatatttgtatcgtcacaccacaattgcatatcaacgatgttagagacatcaagtcattcaacgatatcagattgtactcagaatcccctattaaaacattttccgttattgataacactatcacatatttgttttacttcaaggaacatttgggtgttatttttaacatcactaagtcaagacatgatagaagagtaacaaaattttcaccattgtccaagttcagtgacgtcaaaaatatcactgtttgtttcgatatcgaaagttacttcgacccagaaaaggaatctaaccaagttaacatccctttcatttgttgcgcttcaatcatctataataaggttattggtaacatcgtcgattttgaaggtagagactgtgtcgcacaaatgatcgaatacgttgtagatatttgcggtgaattgaacataagttctgttgaattgatcgcccataacggtggtggttatgatttccactacatcttgtcatccatgtataatccagctgcaattaaaaacatattgatcagaaacaactcttttatttcttttaatttcgcacatgatggtgtcaagttctctgttaaggacagttactcatttttgttgtgtagtgttgccaacgcttctaaagcatttttaaacgaagaaacttttaagaaaactgatttccctcatcacgacttgaaaactgccgatgacttatataaggtctacaaggaatggagttctgttaacacagaaattaaccatgtcgttgaaaaggaaaagttgttgatcacctctgaacacatcgttaacttcaccaagaacgataagagtaagactttgatcgaatggtctaaggattactgtagaaacgacgtattggtcttgtctaaggtttggttggagtttaaaaacgctgtagaagatatttttaactgcgaattggtagaccaaaccatgactttggctggtttgtcatacaagttgttccaagcaaacatgccattcgatgtcgaattgagacatcctaataaggaagactacttcaacatgagagaagcattgattggtggtagatgtatctctgttaacggtatctacaaggatgttttgtgctatgacgtaaaatcattatatccagcttctatggctttctatgatcaaccttacggttcttttaagagagtttcatccagaccaaaagatgaattgggtatctattacgttagagtaaccccaaacagaaacaacaagtcaaactttttccctatcagatcccataataagatcacttacaacaacttcgaagaatcaacttatattgcttggtacacaaacgttgatatcgacattggtttgtcagaaggtcacaacatcgaatacatccctttcgattcctatggtaacattggttactcatggtccaaaaagggtaaaatttttgaaaagtacatcaaggatgttttgtacaaattgaagataaaatatgaaaagcaaaataacaaggtaaaaagaaatgtcatcaaaaagatcatgaactcattgtggggtaaattcgcccaaaagtgggttaacttcgaatacttcatcaagtcagaagatgacattgatttcgaatccgaagaagcatataaaattaacgatacagacttcatgttgatcaaaaagattaaggaatctacctacagttctaagccaatacaaaacggtgtttttactttaagttgggccagatatcacatgaagtctatttgggatgcaggtgccaaagaaggtgctgaatgtatctattcagatacagactccatattcgttcataaggaacacttcaaaaagaatgcaaagtttatgttgaacggtttgaaagtaccaatcatcggttctgaagtcggtcaattggaattagaatgtgaatttgataaattgttgtgcgccggtaaaaagcaatacatgggtttctacacttacttccaagatggtaaaccttgcatcaaggaaaagaaaagattcaaaggtataccaagtaactacatcatacctgaattgtatgctcacttgttatctggtgcagataaggaagccaaaattcaatttttaaaattcagaagagaatggggttccgtcaagggttacatagaaaataagactgtcaaagccacctaactcgaggcgaatttcttatgatttatgatttttattattaaataagttataaaaaaaataagtgtatacaaattttaaagtgactcttaggttttaaaacgaaaattcttattcttgagtaactctttcctgtaggtcaggttgctttctcaggtatagcatgaggtcgctcactagtccatggacgcgtggtaccggccgcgttgctggcgtttttccataggctcggcccccctgacgagcatcacaaaaatcgacgctcaagtcagaggtggcgaaacccgacaggactataaagataccaggcgtttccccctggaagctccctcgtgcgctctcctgttccgaccctgccgcttaccggatacctgtccgcctttctcccttcgggaagcgtggcgctttctcaatgctcacgctgtaggtatctcagttcggtgtaggtcgttcgctccaagctgggctgtgtgcacgaaccccccgttcagcccgaccgctgcgccttatccggtaactatcgtcttgagtccaacccggtaagacacgacttatcgccactggcagcagccactggtaacaggattagcagagcgaggtatgtaggcggtgctacagagttcttgaagtggtggcctaactacggctacactagaaggacagtatttggtatctgcgctctgctgaagccagttaccttcggaaaaagagttggtagctcttgatccggcaaacaaaccaccgctggtagcggtggtttttttgtttgcaagcagcagattacgcgcagaaaaaaaggatctcaagaagatcctttgatcttttctacggggtctgacgctcagtggaacgaaaactcacgttaagggattttggtcatgactgatccttcaactcagcaaaagttcgatttattcaacaaagccacgttgtgtctcaaaatctctgatgttacattgcacaagataaaaatatatcatcatgaacaataaaactgtctgcttacataaacagtaatacaaggggtgttatgagccatattcaacgggaaacgtcttgctccaggccgcgattaaattccaacatggatgctgatttatatgggtataaatgggctcgcgataatgtcgggcaatcaggtgcgacaatctatcgattgtatgggaagcccgatgcgccagagttgtttctgaaacatggcaaaggtagcgttgccaatgatgttacagatgagatggtcagactaaactggctgacggaatttatgcctcttccgaccatcaagcattttatccgtactcctgatgatgcatggttactcaccactgcgatccccgggaaaacagcattccaggtattagaagaatatcctgattcaggtgaaaatattgttgatgcgctggcagtgttcctgcgccggttgcattcgattcctgtttgtaattgtccttttaacagcgatcgcgtatttcgtctcgctcaggcgcaatcacgaatgaataacggtttggttgatgcgagtgattttgatgacgagcgtaatggctggcctgttgaacaagtctggaaagaaatgcataagcttttgccattctcaccggattcagtcgtcactcatggtgatttctcacttgataaccttatttttgacgaggggaaattaataggttgtattgatgttggacgagtcggaatcgcagaccgataccaggatcttgccatcctatggaactgcctcggtgagttttctccttcattacagaaacggctttttcaaaaatatggtattgataatcctgatatgaataaattgcagtttcatttgatgctcgatgagtttttctaatcagaattggttaattggttgtaacacggtccttttcatcacgtgctataaaaataattataatttaaatttttaatataaatatataaattaaaaatagaaagtaaaaaaagaaattaaagaaaaaatagtttttgttttccgaagatgtaaaagactctagggggatcgccaacaaatactaccttttatcttgctcttcctgctctcaggtattaatgccgaattgtttcatcttgtctgtgtagaagaccacacacgaaaatcctgtgattttacattttacttatcgttaatcgaatgtatatctatttaatctgcttttcttgtctaataaatatatatgtaaagtacgctttttgttgaaattttttaaacctttgtttatttttttttcttcattccgtaactcttctaccttctttatttactttctaaaatccaaatacaaaacataaaaataaataaacacagagtaaattcccaaattattccatcattaaaagatacgaggcgcgtgtaagttacaggcaagcgatccgtcctaagaaaccattattatcatgacattaacctataaaaataggcgtatcacgaggccctttcgtctcgcgcgtttcggtgatgacggtgaaaacctctgacacatgcagctcccggagacggtcacagcttgtctgtaagcggatgccgggagcagacaagcccgtcagggcgcgtcagcgggtgttggcgggtgtcggggctggcttaactatgcggcatcagagcagattgtactgagagtgcaccacgcttttcaattcaattcatcattttttttttattcttttttttgatttcgaccgtcgaggagaacttctagtatatctacatacctaatattattgccttattaaaaatggaatcccaacaattacatcaaaatccacattctcttcaaaatcaattgtcctgtacttccttgttcatgtgtgttcaaaaacgttatatttataggataattatactctatttctcaacaagtaattggttgtttggccgagcggtctaaggcgcctgattcaagaaatatcttgaccgcagttaactgtgggaatactcaggtatcgtaagatgcaagagttcgaatctcttagcaaccattatttttttcctcaacataacgagaacacacaggggcgctatcgcacagaatcaaattcgatgactggaaattttttgttaatttcagaggtcgcctgacgcatatacctttttcaactgaaaaattgggagaaaaaggaaaggtgagagcgccggaaccggcttttcatatagaatagagaagcgttcatgactaaatgcttgcatcacaatacttgaagttgacaatattatttaaggacctattgttttttccaataggtggttagcaatcgtcttactttctaacttttcttaccttttacatttcagcaatatatatatatatatttcaaggatataccattctaatgtctgcccctaagaagatcgtcgttttgccaggtgaccacgttggtcaagaaatcacagccgaagccattaaggttcttaaagctatttctgatgttcgttccaatgtcaagttcgatttcgaaaatcatttaattggtggtgctgctatcgatgctacaggtgttccacttccagatgaggcgctggaagcctccaagaaggctgatgccgttttgttaggtgctgtgggtggtcctaaatggggtaccggtagtgttagacctgaacaaggtttactaaaaatccgtaaagaacttcaattgtacgccaacttaagaccatgtaactttgcatccgactctcttttagacttatctccaatcaagccacaatttgctaaaggtactgacttcgttgttgtcagagaattagtgggaggtatttactttggtaagagaaaggaagacgatggtgatggtgtcgcttgggatagtgaacaatacaccgttccagaagtgcaaagaatcacaagaatggccgctttcatggccctacaacatgagccaccattgcctatttggtccttggataaagctaatgttttggcctcttcaagattatggagaaaaactgtggaggaaaccatcaagaacgaatttcctacattgaaggttcaacatcaattgattgattctgccgccatgatcctagttaagaacccaacccacctaaatggtattataatcaccagcaacatgtttggtgatatcatctccgatgaagcctccgttatcccaggttccttgggtttgttgccatctgcgtccttggcctctttgccagacaagaacaccgcatttggtttgtacgaaccatgccacggttctgctccagatttgccaaagaataaggtcaaccctatcgccactatcttgtctgctgcaatgatgttgaaattgtcattgaacttgcctgaagaaggtaaggccattgaagatgcagttaaaaaggttttggatgcaggtatcagaactggtgatttaggtggttccaacagtaccaccgaagtcggtgatgctgtcgccgaagaagttaagaaaatccttgcttaaaaagattctctttttttatgatatttgtacataaactttataaatgaaattcataatagaaacgacacgaaattacaaaatggaatatgttcatagggtagacgaaactatatacgcaatctacatacatttatcaagaaggagaaaaaggaggatgtaaaggaatacaggtaagcaaattgatactaatggctcaacgtgataaggaaaaagaattgcactttaacattaatattgacaaggaggagggcaccacacaaaaagttaggtgtaacagaaaatcatgaaactatgattcctaatttatatattggaggattttctctaaaaaaaaaaaaaatacaacaaataaaaaacactcaatgacctgaccatttgatggagtttaagtcaataccttcttgaaccatttcccataatggtgaaagttccctcaagaattttactctgtcagaaacggccttaacgacgtagtcgatatgcggtgtgaaataccgcacagatgcgtaaggagaaaataccgcatcagg |
| pAW729 | CEN/ARS plasmid encoding BadBoy3 DNA polymerase with a LEU2 marker for yeast and a KanR marker for propagation in *E. coli*. | gagctcgtgttgttatccgatacaaccggatatttttcttttaatgagtctaaaccgtgatagcttcaggttaatacaatcaaaaaaagctcaaatattcttttaatgccgcgttcacagattccaattgaatacaactaggtagttcattatatgaagcctttgctactatttttcactatagtctgccttcaccttaatgcagacatccacatattttaatcactttaaaataaaaaggaagatatattagaagctatgatccaatctgtaagccagattaaaattcacgaactcttctttcatttgaattgaatgctttgagttggggtagattatcgcaaattactcatcacatttattgactacgaacttgctgatgtcctttttttatttatatttttcttcagtgaagcgatttttttttacacagaccaagacggaaaaaagtagctaaggaagaaaacaaaatcatgaaaaaaatgtgaagtgatcatgcacatcgcatcaacttaaacattggcttagagatatatagagttagagtttacggcaacctttaagcaccaatacctttttggcatagtctaaagacctggttcttaattttaaacaaatttaactaaagatttccctatcaaagaagtaacgagttgacagattttctcaaaataaatcgatactgcatttctaggcatatccagcgagatctatggactacaaagacaaagcattaaacgacttgagaaacgtatacgcagacttcgactccttgccattggacttcagacaaatcttaattaaggatagagctacattgttgcaaaaggaagacgttgaaaagaaaattttggaaagacaagaagatgccaaaaagtacgctgaatacttgaagcaatcagaaatcccagaaagaatctccttacctaacatcaagagacataagggtgtatcaatctcctttgaagaaacatctgaagatatggtcttggaaccaagaccttttatcttcgacggtttgaacatcagatgtttcagaagagaaaccatattcagtttgaaaaataagattttgaacatggttaaggaatcttcatcctttaaaaacgtttcaagacaaagtgtatcttttatgtacttcaaaatttttaataagggtaaagttattgcaagtacaaagtctgttaacatctatgaagataagatcgacgaaagattggaagatttgtgtaacaacttcgatgacgtattgaaaaagattatagacgtcacctatggttacgaatccttattcgttagtgaaacttactcttacgtaatattctatgctaaatcaatctactttccacaacctagatgcgttaacaactggggtaacaacatcccaaacatcttgacattcgattcctttaagcttttcaccgcaaacaaaaacaacgttagttgtatcaagcaatgttcaagatttttgtggcaaaaagatttcaacactttagaagaaatgatagaatacaagaacggtaatatttgtatcgtcacaccacaattgcatatcagcgatgttagagacatcaagtcattcaacgatatcagattgtactcagaatcccctattaaaacattttccgttattgataacactatcacatatttgttttacttcaaggaacatttgggtgttatttttaacttcactaagtcaagacatgatagaagagtaacaaaattttcaccattgtccaagttcagtgacgtcaaaaatatcactgtttgtttcgatatcgaaagttacttcgacccagaaaaggaatctaaccaagttaacatccctttcatttgttgcgcttcaatcatctataataaggttattggtaacatcgtcgattttgaaggtagagactgtgtcgcacaaatgatcgaatacgttgtagatatttgcggtgaattgaacataagttctgttgaattgatcgcccataacggtggtggttatgatttccactacatcttgtcatccatgtataatccagctgcaattaaaaacatattgatcagaaacaactcttttatttcttttaatttcgcacatgatggtgtcaagttctctgttaaggacagttactcatttttgtcgtgtagtttggccaacgcttctaaagcatttttaaacgaaggaacttttaagaaaactgatttccctcatcacgacttgaaaactgccgatgacttatataaggtctacaaggaatggagttctgttaacacagaaattaaccatgtcgttgaaaaggaaaagttgttgatcacctctgaacacatcgttaacttcaccaagaacgataagagtaagactttgatcgaatggtctaaggattactgtagaaacgacgtattggtcttgtctaaggtttggttggagtttaaaaacgctgtagaagatatttttaactgcgaattggtagaccaaaccatgactttggctggtttgtcatacaagttgttccaagcaaacatgccattcgatgtcgaattgagacatcctaataaggaagactacttcaacatgagagaagcattgattggtggtagatgtatctctgttaacggtatctacagggatgttttgtgcttagacgtaaaatcattatatccagcttctatggctttctatgatcaaccttacggttcttttaagagagtttcatccagaccaaaagatgaattgggtatctattacgttagagtaaccacaaacagaaacaacaagtcaaactttttccctatcagatcccataataagatcacttacaacaacttcgaagaatcaacttatattgcttggtacacaaacgttgatattgacattggtttgtcagaaggtcacaacatcgaatacatccctttcgattcctatggtaacattggttactcatggtccaaaaagggtaaaatttttgaaaagtacatcagggatgttttgtacaaattgaagataaaatatgaaaagcaaaataacaaggtaaaaagaaatgtcatcaaaaagatcatgaactcattgtggggtaaattcgcccaaaagtgggttaacttcgaatacttcatcaagtcagaagatgacattgatttcgaatccgaagaagcatataaaatttgggatacagacttcatgttgatcaaaaagattaaggaatctatctacagttctaagccaatacaaaacggtgtttttactttaagttgggccagatatcacatgaagtctatttgggatgcaggtgccaaagaaggtgctgaatgtatctattcagatacagactccatattcgttcataaggaacacttcaaaaagaatgcaaagtttatgttgaacggtttgaaagtaccaatcatcggttctgaagtcggtcaatctgaattagaatgtgaatttgataaattgttgtgcgccggtaaaaagcaatacatgggtttctacacttacttccaagatggtaaaccttgcatcaaggaaaagaaaagattcaaaggtataccaagtaactacatcatacctgaattgtatgctcacttgttatctggtgcagataaggaagccaaaattcaatatttaaaattcagaagagaatggggttccgtcaagggttacatagaaaataagactgtcaaagccacctaactcgaggcgaatttcttatgatttatgatttttattattaaataagttataaaaaaaataagtgtatacaaattttaaagtgactcttaggttttaaaacgaaaattcttattcttgagtaactctttcctgtaggtcaggttgctttctcaggtatagcatgaggtcgctcactagtccatggacgcgtggtaccggccgcgttgctggcgtttttccataggctcggcccccctgacgagcatcacaaaaatcgacgctcaagtcagaggtggcgaaacccgacaggactataaagataccaggcgtttccccctggaagctccctcgtgcgctctcctgttccgaccctgccgcttaccggatacctgtccgcctttctcccttcgggaagcgtggcgctttctcaatgctcacgctgtaggtatctcagttcggtgtaggtcgttcgctccaagctgggctgtgtgcacgaaccccccgttcagcccgaccgctgcgccttatccggtaactatcgtcttgagtccaacccggtaagacacgacttatcgccactggcagcagccactggtaacaggattagcagagcgaggtatgtaggcggtgctacagagttcttgaagtggtggcctaactacggctacactagaaggacagtatttggtatctgcgctctgctgaagccagttaccttcggaaaaagagttggtagctcttgatccggcaaacaaaccaccgctggtagcggtggtttttttgtttgcaagcagcagattacgcgcagaaaaaaaggatctcaagaagatcctttgatcttttctacggggtctgacgctcagtggaacgaaaactcacgttaagggattttggtcatgactgatccttcaactcagcaaaagttcgatttattcaacaaagccacgttgtgtctcaaaatctctgatgttacattgcacaagataaaaatatatcatcatgaacaataaaactgtctgcttacataaacagtaatacaaggggtgttatgagccatattcaacgggaaacgtcttgctccaggccgcgattaaattccaacatggatgctgatttatatgggtataaatgggctcg |

| | | |
|---|---|---|
| | | cgataatgtcgggcaatcaggtgcgacaatctatcgattgtatgggaagcccgatgcgccagagttgtttctgaaacatggcaaaggtagcgttgccaatgatgttacagatgagatggtcagac<br>taaactggctgacggaatttatgcctcttccgaccatcaagcattttatccgtactcctgatgatgcatggttactcaccactgcgatccccgggaaaacagcattccaggtattagaagaatatcct<br>gattcaggtgaaaatattgttgatgcgctggcagtgttcctgcgccggttgcattcgattcctgtttgtaattgtccttttaacagcgatcgcgtatttcgtctcgctcaggcgcaatcacgaatgaata<br>acggtttggttgatgcgagtgattttgatgacgagcgtaatggctggcctgttgaacaagtctggaaagaaatgcataagcttttgccattctcaccggattcagtcgtcactcatggtgatttctcac<br>ttgataaccttatttttgacgaggggaaattaataggttgtattgatgttggacgagtcggaatcgcagaccgataccaggatcttgccatcctatggaactgcctcggtgagttttctccttcattaca<br>gaaacggctttttcaaaaatatggtattgataatcctgatatgaataaattgcagtttcatttgatgctcgatgagtttttctaatcagaattggttaattggttgtaacacggtccttttcatcacgtgctat<br>aaaaataattataatttaaatttttttaatataaatatataaattaaaaatagaaagtaaaaaaagaaattaaagaaaaaatagtttttgttttccgaagatgtaaaagactctaggggggatcgccaacaaa<br>tactaccttttatcttgctcttcctgctctcaggtattaatgccgaattgtttcatcttgtctgtgtagaagaccacacacgaaaatcctgtgattttacattttacttatcgttaatcgaatgtatatctatttaa<br>tctgcttttcttgtctaataaatatatatgtaaagtacgctttttgttgaaattttttaaacctttgtttattttttttttcttcattccgtaactcttctaccttctttatttactttctaaaatccaaatacaaaacataaa<br>aataaataaacacagagtaaattcccaaattattccatcattaaaagatacgaggcgcgtgtaagttacaggcaagcgatccgtcctaagaaaccattattatcatgacattaacctataaaaatag<br>gcgtatcacgaggccctttcgtctcgcgcgtttcggtgatgacggtgaaaacctctgacacatgcagctcccggagacggtcacagcttgtctgtaagcggatgccgggagcagacaagccc<br>gtcagggcgcgtcagcgggtgttggcgggtgtcggggctggcttaactatgcggcatcagagcagattgtactgagagtgcaccacgcttttcaattcaattcatcatttttttttttattcttttttttgat<br>ttcgaccgtcgaggagaacttctagtatatctacatacctaatattattgccttattaaaaatggaatcccaacaattacatcaaaatccacattctcttcaaaatcaattgtcctgtacttccttgttcatgt<br>gtgttcaaaaacgttatatttataggataattatactctatttctcaacaagtaattggttgtttggccgagcggtctaaggcgcctgattcaagaaatatcttgaccgcagttaactgtgggaatactca<br>ggtatcgtaagatgcaagagttcgaatctcttagcaaccattatttttttcctcaacataacgagaacacacaggggcgctatcgcacagaatcaaattcgatgactggaaattttttgttaatttcaga<br>ggtcgcctgacgcatataccttttttcaactgaaaaattgggagaaaaaggaaaggtgagagcgccggaaccggcttttcatatagaatagagaagcgttcatgactaaatgcttgcatcacaata<br>cttgaagttgacaatattatttaaggacctattgttttttccaataggtggttagcaatcgtcttactttctaacttttcttaccttttacatttcagcaatatatatatatatatttcaaggatataccattctaatg<br>tctgcccctaagaagatcgtcgttttgccaggtgaccacgttggtcaagaaatcacagccgaagccattaaggttcttaaagctatttctgatgttcgttccaatgtcaagttcgatttcgaaaatcatt<br>taattggtggtgctgctatcgatgctacaggtgttccacttccagatgaggcgctggaagcctccaagaaggctgatgccgttttgttaggtgctgtgggtggtcctaaatggggtaccggtagtgt<br>tagacctgaacaaggtttactaaaaatccgtaaagaacttcaattgtacgccaacttaagaccatgtaactttgcatccgactctcttttagacttatctccaatcaagccacaatttgctaaaggtact<br>gacttcgttgttgtcagagaattagtgggaggtatttactttggtaagagaaaggaagacgatggtgatggtgtcgcttgggatagtgaacaatacaccgttccagaagtgcaaagaatcacaag<br>aatggccgctttcatggccctacaacatgagccaccattgcctatttggtccttggataaagctaatgttttggcctcttcaagattatggagaaaaactgtggaggaaaccatcaagaacgaatttc<br>ctacattgaaggttcaacatcaattgattgattctgccgccatgatcctagttaagaacccaacccacctaaatggtattataatcaccagcaacatgtttggtgatatcatctccgatgaagcctccg<br>ttatcccaggttccttgggtttgttgccatctgcgtccttggcctctttgccagacaagaacaccgcatttggtttgtacgaaccatgccacggttctgctccagatttgccaaagaataaggtcaacc<br>ctatcgccactatcttgtctgctgcaatgatgttgaaattgtcattgaacttgcctgaagaaggtaaggccattgaagatgcagttaaaaaggttttggatgcaggtatcagaactggtgatttaggtg<br>gttccaacagtaccaccgaagtcggtgatgctgtcgccgaagaagttaagaaaatccttgcttaaaaagattctctttttttatgatatttgtacataaactttataaatgaaattcataatagaaacgac<br>acgaaattacaaaatggaatatgttcatagggtagacgaaactatatacgcaatctacatacatttatcaagaaggagaaaaaggaggatgtaaaggaatacaggtaagcaaattgatactaatg<br>gctcaacgtgataaggaaaaagaattgcactttaacattaatattgacaaggaggagggcaccacacaaaaagttaggtgtaacagaaaatcatgaaactatgattcctaatttatatattggagga<br>ttttctctaaaaaaaaaaaaatacaacaaataaaaaacactcaatgacctgaccatttgatggagtttaagtcaataccttcttgaaccatttcccataatggtgaaagttccctcaagaattttactctg<br>tcagaaacggccttaacgacgtagtcgatatgcggtgtgaaataccgcacagatgcgtaaggagaaaataccgcatcagg |
| pAP175 | CEN/ARS plasmid for galactose-inducible yeast surface display with LEU2 selection marker for yeast and an AmpR marker for propagation in *E. coli*. | tcgaggagaacttctagtatatctacatacctaatattattgccttattaaaaatggaatcccaacaattacatcaaaatccacattctcttcaaaatcaattgtcctgtacttccttgttcatgtgtgttcaa<br>aaacgttatatttataggataattatactctatttctcaacaagtaattggttgtttggccgagcggtctaaggcgcctgattcaagaaatatcttgaccgcagttaactgtgggaatactcaggtatcgt<br>aagatgcaagagttcgaatctcttagcaaccattatttttttcctcaacataacgagaacacacaggggcgctatcgcacagaatcaaattcgatgactggaaattttttgttaatttcagaggtcgcc<br>tgacgcatataccttttttcaactgaaaaattgggagaaaaaggaaaggtgagagcgccggaaccggcttttcatatagaatagagaagcgttcatgactaaatgcttgcatcacaatacttgaagtt<br>gacaatattatttaaggacctattgttttttccaataggtggttagcaatcgtcttactttctaacttttcttaccttttacatttcagcaatatatatatatatatttcaaggatataccattctaatgtctgcccc<br>taagaagatcgtcgttttgccaggtgaccacgttggtcaagaaatcacagccgaagccattaaggttcttaaagctatttctgatgttcgttccaatgtcaagttcgatttcgaaaatcatttaattggt<br>ggtgctgctatcgatgctacaggtgttccacttccagatgaggcgctggaagcctccaagaaggctgatgccgttttgttaggtgctgtgggtggtcctaaatggggaaccggtagtgttagacct<br>gaacaaggtttactaaaaatccgtaaagaacttcaattgtacgccaacttaagaccatgtaactttgcatccgactctcttttagacttatctccaatcaagccacaatttgctaaaggtactgacttcgt<br>tgttgtcagagaattagtgggaggtatttactttggtaagagaaaggaggacgatggtgatggtgtcgcttgggatagtgaacaatacaccgttccagaagtgcaaagaatcacaagaatggcc<br>gctttcatggccctacaacatgagccaccattgcctatttggtccttggataaagctaatgttttggcctcttcaagattatggagaaaaactgtggaggaaaccatcaagaacgaatttcctacattg<br>aaggttcaacatcaattgattgattctgccgccatgatcctagttaagaacccaacccacctaaatggtattataatcaccagcaacatgtttggtgatatcatctccgatgaagcctccgttatccca<br>ggttccttgggtttgttgccatctgcgtccttggcctctttgccagacaagaacaccgcatttggtttgtacgaaccatgccacggttctgctccagatttgccaaagaataaggtcaaccctatcgc<br>cactatcttgtctgctgcaatgatgttgaaattgtcattgaacttgcctgaagaaggtaaggccattgaagatgcagttaaaaaggttttggatgcaggtatcagaactggtgatttaggtggttccaa<br>cagtaccaccgaagtcggtgatgctgtcgccgaagaagttaagaaaatccttgcttaacctggaggacccttctctttagactattctactcttatgcacgtaaaaaattctaggaaatatgtattaac<br>taggagtaaaataaccggctagtggcattcatatagccgtctgtttacatctacatcacacatttcgagtgtatatctcgcaacgttggcgttaaataggcagatcacgtgctataaaaataattataat<br>ttaaatttttttaatataaatatataaattaaaaatagaaagtaaaaaaagaaattaaagaaaaaatagtttttgttttccgaagatgtaaaagactctaggggggatcgccaacaaatactaccttttatctt<br>gctcttcctgctctcaggtattaatgccgaattgtttcatcttgtctgtgtagaaaaccacacacgaaaatcctgtgattttacattttacttatcgttaatcgaatgtatatctatttaatctgcttttcttgtct<br>aataaatatatatgtaaagtacgctttttgttgaaattttttaaacctttgtttattttttttttcttcattccgtaactcttctaccttctttatttactttctaaaatccaaatacaaaacataaaaataaataaacac<br>agagtaaattcccaaattattccatcattaaaagatacgaggcgcgtgtaagttacaggcaagcgatccgtcccgagcggccgcgattatcaaaaaggatcttcacctagatccttttaaattaaaa<br>atgaagttttaaatcaatctaaagtatatatgagtaaacttggtctgacagttaccaatgcttaatcagtgaggcacctatctcagcgatctgtctatttcgttcatccatagttgcctgactccccgtcgt<br>gtagataactacgatacgggagggcttaccatctggccccagtgctgcaatgataccgcgggacccacgctcaccggctccagatttatcagcaataaaccagccagccggaagggccgag<br>cgcagaagtggtcctgcaactttatccgcctccatccagtctattaattgttgccgggaagctagagtaagtagttcgccagttaatagtttgcgcaacgttgttgccattgctacaggcatcgtggt<br>gtcacgctcgtcgtttggtatggcttcattcagctccggttcccaacgatcaaggcgagttacatgatcccccatgttgtgcaaaaaagcggttagctccttcggtcctccgatcgttgtcagaagta<br>agttggccgcagtgttatcactcatggttatggcagcactgcataattctcttactgtcatgccatccgtaagatgcttttctgtgactggtgagtactcaaccaagtcattctgagaatagtgtatgcg<br>gcgaccgagttgctcttgcccggcgtcaatacgggataataccgcgccacatagcagaactttaaaagtgctcatcattggaaaacgttcttcggggcgaaaactctcaaggatcttaccgctgtt<br>gagatccagttcgatgtaacccactcgtgcacccaactgatcttcagcatcttttactttcaccagcgtttctgggtgagcaaaaacaggaaggcaaaatgccgcaaaaaagggaataagggcg<br>acacggaaatgttgaatactcatactcttcctttttcaatattattgaagcatttatcagggttattgtctcatgagcggatacatatttgaatgtatttagaaaaataaacaaataggggttccgcgcaca<br>tttccccgaaaagtgccacctgtcatgaccaaaatcccttaacgtgagttttcgttccactgagcgtcagaccccgtagaaaagatcaaaggatcttcttgagatcctttttttctgcgcgtaatctgct<br>gcttgcaaacaaaaaaaccaccgctaccagcggtggtttgtttgccggatcaagagctaccaactctttttccgaaggtaactggcttcagcagagcgcagataccaaatactgttcttctagtgta<br>gccgtagttaggccaccacttcaagaactctgtagcaccgcctacatacctcgctctgctaatcctgttaccagtggctgctgccagtggcgataagtcgtgtcttaccgggttggactcaagacg<br>atagttaccggataaggcgcagcggtcgggctgaacggggggttcgtgcacacagcccagcttggagcgaacgacctacaccgaactgagatacctacagcgtgagctatgagaaagcgc<br>cacgcttcccgaagggagaaaggcggacaggtatccggtaagcggcagggtcggaacaggagagcgcacgagggagcttccagggggaaacgcctggtatctttatagtcctgtcgggtt<br>tcgccacctctgacttgagcgtcgatttttgtgatgctcgtcaggggggcggagcctatggaaaaacgccagcaacgcggcctttttacggttcctggccttttgctggccttttgctcacatgttctt<br>tcctgcgttatcccctgattctgtggataaccgtgcggccgcccctgaattcgcatctagactgatgagacgtggtagagccacaaacagccggtacaagcaacgatctccaggaccatctgaat<br>catgcgcggatgacacgaactcacgacggcgatcacagacattaacccacagtacagacactgcgacaacgtggcaattcgtcgcaatacaacgcctaggccccattatcttagcctaaaaaa<br>accttctctttggaactttcagtaatacgcttaactgctcattgctatattgaagtacggattagaagccgccgagcgggcgacagccctccgacggatgactctcctccgtgcgtcctcgtcatca<br>ccggtcgcgttcctgaaacgcagatgtgcctcgcgccgcactgctccgaacaataaagattctacaatactagcttttatggttatgaagaggaaaaattggcagtaacctggccccacaaacctt<br>caaattaacgaatcaaattaacaaccataggatgataatgcgattagttttttagccttatttctggggtaattaatcagcgaagcgatgatttttgatctattaacagatatataaatggaaaagctgca<br>taaccactttaactaatactttcaacattttcagtttgtattacttcttattcaaatgtcataaaagtatcaacaaaaaattgttaatatacctctatactttaacgtcaaggagaaaaaactataatgagattc<br>ccatctatcttcaccgctgttttgttcgctgcttcttctgctttggctgctccagctaacaccaccaccgaagacgaaaccgctcaaatcccagctgaagctgttatcgactactctgacttggaaggt<br>gacttcgacgctgctgctttgccattgtctaactctaccaacaacggtttgtcttctaccaacaccaccatcgcttctatcgctgctaaggaagaaggtgttcaattggacaagagagaagctgacg<br>caggtaccgagaccgaaagtgaaacgtgatttcatgcgtcattttgaacattttgtaaatcttatttaataatgtgtgcggcaattcacatttaatttatgaatgttttcttaacatcgcggcaactcaaga<br>aacggcaggttcggatcttagctactagagaaagaggagaaatactagatgcgtaaaggcgaagagctgttcactggtgtcgtccctattctggtggaactggatggtgatgtcaacggtcataa<br>gttttccgtgcgtggcgagggtgaaggtgacgcaactaatggtaaactgacgctgaagttcatctgtactactggtaaactgccggttccttggccgactctggtaacgacgctgacttatggtgtt<br>cagtgctttgctcgttatccggaccatatgaagcagcatgacttcttcaagtccgccatgccggaaggctatgtgcaggaacgcacgatttcctttaaggatgacggcacgtacaaaacgcgtgc<br>ggaagtgaaatttgaaggcgataccctggtaaaccgcattgagctgaaaggcattgactttaaagaggacggcaatatcctgggccataagctggaatacaattttaacagccacaatgtttacat<br>caccgccgataaacaaaaaaatggcattaaagcgaattttaaaattcgccacaacgtggaggatggcagcgtgcagctggctgatcactaccagcaaaacactccaatcggtgatggtcctgtt<br>ctgctgccagacaatcactatctgagcacgcaaagcgttctgtctaaagatccgaacgagaaacgcgatcatatggttctgctggagttcgtaaccgcagcgggcatcacgcatggtatggatg<br>aactgtacaaatgaccaggcatcaaataaaacgaaaggctcagtcgaaagactgggcctttcgttttatctgttgtttgtcggtgaacgctctctactagagtcacactggctcaccttcgggtggg<br>cctttctgcgtttataggtctcgtcgacggttccaaggacaatagctcgacgattgaaggtagatacccatacgacgttccagactacgctctgcaggctagtggtggtggtggttctggtggtggt<br>ggttctggtggtggtggttctgctagcgacgtccaggaactgacaactatatgcgagcaaatcccctcaccaactttagaatcgacgccgtactctttgtcaacgactactattttggccaacggga<br>aggcaatgcaaggagtttttgaatattacaaatcagtaacgtttgtcagtaattgcggttctcacccctcaacaactagcaaaggcagccccataaacacacagtatgtttttgatagctcgagatct<br>gataacaacagtgtagatgtaacaaaatcgactttgttcccactgtacttttagctcgtacaaaatacaatatacttttcatttctccgtaaacaacatgttttcccatgtaatatccttttctatttttcgttcc<br>gttaccaactttacacatactttatatagctattcacttctatacactaaaaaactaagacaattttaattttgctgcctgccatatttcaatttgttataaattcctataatttatcctattagtagctaaaaaa<br>gatgaatgtgaatcgaatcctaagagaatt |
| pAP176 | CEN/ARS plasmid for β-estradiol-inducible yeast surface display with LEU2 selection marker for yeast and an AmpR marker for propagation in *E. coli*. | tcgaggagaacttctagtatatctacatacctaatattattgccttattaaaaatggaatcccaacaattacatcaaaatccacattctcttcaaaatcaattgtcctgtacttccttgttcatgtgtgttcaa<br>aaacgttatatttataggataattatactctatttctcaacaagtaattggttgtttggccgagcggtctaaggcgcctgattcaagaaatatcttgaccgcagttaactgtgggaatactcaggtatcgt<br>aagatgcaagagttcgaatctcttagcaaccattatttttttcctcaacataacgagaacacacaggggcgctatcgcacagaatcaaattcgatgactggaaattttttgttaatttcagaggtcgcc<br>tgacgcatataccttttttcaactgaaaaattgggagaaaaaggaaaggtgagagcgccggaaccggcttttcatatagaatagagaagcgttcatgactaaatgcttgcatcacaatacttgaagtt<br>gacaatattatttaaggacctattgttttttccaataggtggttagcaatcgtcttactttctaacttttcttaccttttacatttcagcaatatatatatatatatttcaaggatataccattctaatgtctgcccc<br>taagaagatcgtcgttttgccaggtgaccacgttggtcaagaaatcacagccgaagccattaaggttcttaaagctatttctgatgttcgttccaatgtcaagttcgatttcgaaaatcatttaattggt<br>ggtgctgctatcgatgctacaggtgttccacttccagatgaggcgctggaagcctccaagaaggctgatgccgttttgttaggtgctgtgggtggtcctaaatggggaaccggtagtgttagacct<br>gaacaaggtttactaaaaatccgtaaagaacttcaattgtacgccaacttaagaccatgtaactttgcatccgactctcttttagacttatctccaatcaagccacaatttgctaaaggtactgacttcgt<br>tgttgtcagagaattagtgggaggtatttactttggtaagagaaaggaggacgatggtgatggtgtcgcttgggatagtgaacaatacaccgttccagaagtgcaaagaatcacaagaatggcc<br>gctttcatggccctacaacatgagccaccattgcctatttggtccttggataaagctaatgttttggcctcttcaagattatggagaaaaactgtggaggaaaccatcaagaacgaatttcctacattg<br>aaggttcaacatcaattgattgattctgccgccatgatcctagttaagaacccaacccacctaaatggtattataatcaccagcaacatgtttggtgatatcatctccgatgaagcctccgttatccca<br>ggttccttgggtttgttgccatctgcgtccttggcctctttgccagacaagaacaccgcatttggtttgtacgaaccatgccacggttctgctccagatttgccaaagaataaggtcaaccctatcgc<br>cactatcttgtctgctgcaatgatgttgaaattgtcattgaacttgcctgaagaaggtaaggccattgaagatgcagttaaaaaggttttggatgcaggtatcagaactggtgatttaggtggttccaa<br>cagtaccaccgaagtcggtgatgctgtcgccgaagaagttaagaaaatccttgcttaacctggaggacccttctctttagactattctactcttatgcacgtaaaaaattctaggaaatatgtattaac<br>taggagtaaaataaccggctagtggcattcatatagccgtctgtttacatctacatcacacatttcgagtgtatatctcgcaacgttggcgttaaataggcagatcacgtgctataaaaataattataat<br>ttaaatttttttaatataaatatataaattaaaaatagaaagtaaaaaaagaaattaaagaaaaaatagtttttgttttccgaagatgtaaaagactctaggggggatcgccaacaaatactaccttttatctt<br>gctcttcctgctctcaggtattaatgccgaattgtttcatcttgtctgtgtagaaaaccacacacgaaaatcctgtgattttacattttacttatcgttaatcgaatgtatatctatttaatctgcttttcttgtct<br>aataaatatatatgtaaagtacgctttttgttgaaattttttaaacctttgtttattttttttttcttcattccgtaactcttctaccttctttatttactttctaaaatccaaatacaaaacataaaaataaataaacac<br>agagtaaattcccaaattattccatcattaaaagatacgaggcgcgtgtaagttacaggcaagcgatccgtcccgagcggccgcgattatcaaaaaggatcttcacctagatccttttaaattaaaa<br>atgaagttttaaatcaatctaaagtatatatgagtaaacttggtctgacagttaccaatgcttaatcagtgaggcacctatctcagcgatctgtctatttcgttcatccatagttgcctgactccccgtcgt<br>gtagataactacgatacgggagggcttaccatctggccccagtgctgcaatgataccgcgggacccacgctcaccggctccagatttatcagcaataaaccagccagccggaagggccgag<br>cgcagaagtggtcctgcaactttatccgcctccatccagtctattaattgttgccgggaagctagagtaagtagttcgccagttaatagtttgcgcaacgttgttgccattgctacaggcatcgtggt<br>gtcacgctcgtcgtttggtatggcttcattcagctccggttcccaacgatcaaggcgagttacatgatcccccatgttgtgcaaaaaagcggttagctccttcggtcctccgatcgttgtcagaagta<br>agttggccgcagtgttatcactcatggttatggcagcactgcataattctcttactgtcatgccatccgtaagatgcttttctgtgactggtgagtactcaaccaagtcattctgagaatagtgtatgcg<br>gcgaccgagttgctcttgcccggcgtcaatacgggataataccgcgccacatagcagaactttaaaagtgctcatcattggaaaacgttcttcggggcgaaaactctcaaggatcttaccgctgtt<br>gagatccagttcgatgtaacccactcgtgcacccaactgatcttcagcatcttttactttcaccagcgtttctgggtgagcaaaaacaggaaggcaaaatgccgcaaaaaagggaataagggcg<br>acacggaaatgttgaatactcatactcttcctttttcaatattattgaagcatttatcagggttattgtctcatgagcggatacatatttgaatgtatttagaaaaataaacaaataggggttccgcgcaca |

| | | |
|---|---|---|
| | | tttccccgaaaagtgccacctgtcatgaccaaaatcccttaacgtgagttttcgttccactgagcgtcagaccccgtagaaaagatcaaaggatcttcttgagatcctttttttctgcgcgtaatctgct<br>gcttgcaaacaaaaaaaccaccgctaccagcggtggtttgtttgccggatcaagagctaccaactctttttccgaaggtaactggcttcagcagagcgcagataccaaatactgttcttctagtgta<br>gccgtagttaggccaccacttcaagaactctgtagcaccgcctacatacctcgctctgctaatcctgttaccagtggctgctgccagtggcgataagtcgtgtcttaccgggttggactcaagacg<br>atagttaccggataaggcgcagcggtcgggctgaacggggggttcgtgcacacagcccagcttggagcgaacgacctacaccgaactgagatacctacagcgtgagctatgagaaagcgc<br>cacgcttcccgaagggagaaaggcggacaggtatccggtaagcggcagggtcggaacaggagagcgcacgagggagcttccagggggaaacgcctggtatctttatagtcctgtcgggtt<br>tcgccacctctgacttgagcgtcgatttttgtgatgctcgtcaggggggcggagcctatggaaaaacgccagcaacgcggcctttttacggttcctggccttttgctggccttttgctcacatgttctt<br>tcctgcgttatcccctgattctgtggataaccgtgcggccgcccctgaattcgcatctagactgatgagacgtggtagagccacaaacagccggtacaagcaacgatctccaggaccatctgaat<br>catgcgcggatgacacgaactcacgacggcgatcacagacattaacccacagtacagacactgcgacaacgtggcaattcgtcgcaatacaacgcctaggttgaggccaacgattatggga<br>gatcgaatattaatattatgggagatcgaatattaatattatgggagatcgaatattaatattatgggagatcgaatattaatattatgggagatcgaatattaatattatgggagatcgaatattaatatt<br>atgggagatggcatgaatggccttaagcagatccgccaggcgtgtatatatagcgtggatggccaggcaactttagtgctgacacatacaggcatatatatatgtgtgcgacgacacatgatcat<br>atggcatgcatgtgctctgtatgtatataaaactcttgttttcttcttttctctaaatattctttccttatacattaggacctttgcagcataaattactatacttctatagacacacaaacacaaatacacaca<br>ctaaatgagattcccatctatcttcaccgctgttttgttcgctgcttcttctgctttggctgctccagctaacaccaccaccgaagacgaaaccgctcaaatcccagctgaagctgttatcgactactct<br>gacttggaaggtgacttcgacgctgctgctttgccattgtctaactctaccaacaacggtttgtcttctaccaacaccaccatcgcttctatcgctgctaaggaagaaggtgttcaattggacaagag<br>agaagctgacgcaggtaccgagaccgaaagtgaaacgtgatttcatgcgtcattttgaacattttgtaaatcttatttaataatgtgtgcggcaattcacatttaatttatgaatgttttcttaacatcgcg<br>gcaactcaagaaacggcaggttcggatcttagctactagagaaagaggagaaatactagatgcgtaaaggcgaagagctgttcactggtgtcgtccctattctggtggaactggatggtgatgt<br>caacggtcataagttttccgtgcgtggcgagggtgaaggtgacgcaactaatggtaaactgacgctgaagttcatctgtactactggtaaactgccggttccttggccgactctggtaacgacgct<br>gacttatggtgttcagtgctttgctcgttatccggaccatatgaagcagcatgacttcttcaagtccgccatgccggaaggctatgtgcaggaacgcacgatttcctttaaggatgacggcacgtac<br>aaaacgcgtgcggaagtgaaatttgaaggcgataccctggtaaaccgcattgagctgaaaggcattgactttaaagaggacggcaatatcctgggccataagctggaatacaattttaacagcc<br>acaatgtttacatcaccgccgataaacaaaaaaatggcattaaagcgaattttaaaattcgccacaacgtggaggatggcagcgtgcagctggctgatcactaccagcaaaacactccaatcgg<br>tgatggtcctgttctgctgccagacaatcactatctgagcacgcaaagcgttctgtctaaagatccgaacgagaaacgcgatcatatggttctgctggagttcgtaaccgcagcgggcatcacgc<br>atggtatggatgaactgtacaaatgaccaggcatcaaataaaacgaaaggctcagtcgaaagactgggcctttcgttttatctgttgtttgtcggtgaacgctctctactagagtcacactggctcac<br>cttcgggtgggcctttctgcgtttataggtctcgtcgacggttccaaggacaatagctcgacgattgaaggtagatacccatacgacgttccagactacgctctgcaggctagtggtggtggtggt<br>tctggtggtggtggttctggtggtggtggttctgctagcgacgtccaggaactgacaactatatgcgagcaaatcccctcaccaactttagaatcgacgccgtactctttgtcaacgactactatttt<br>ggccaacgggaaggcaatgcaaggagtttttgaatattacaaatcagtaacgtttgtcagtaattgcggttctcaccccctcaacaactagcaaaggcagccccataaacacacagtatgtttttga<br>tagctcgagatctgataacaacagtgtagatgtaacaaaatcgactttgttcccactgtacttttagctcgtacaaaatacaatatacttttcatttctccgtaaacaacatgttttcccatgtaatatccttt<br>tctatttttcgttccgttaccaactttacacatactttatatagctattcacttctatacactaaaaaactaagacaattttaattttgctgcctgccatatttcaatttgttataaattcctataatttatcctattag<br>tagctaaaaaaagatgaatgtgaatcgaatcctaagagaatt |
| pAP191 | Plasmid for generation of an integration cassette onto p1 where the cassette encodes nanobody-HA-Aga2 fusion with TRP1 and NatMX selection markers. Integration cassette is generated by linearization with ScaI. Contains AmpR marker for propagation in *E. coli*. | gctgcgaatttcttatgatttatgatttttattattaaataagttataaaaaaaataagtgtatacaaattttaaagtgactcttaggttttaaaacgaaaattcttattcttgagtaactctttcctgtaggtcag<br>gttgctttctcaggtatagcatgaggtcgctcgagctcttaagggcaaggcatagacatatacaaggcctgttcaccgtcagatgctgttccatcatataaagcagtatctaaaccacataatgtga<br>aacccattcttctataagcatggatagcaggggcgttaacattagtaacttctaaccataaatgaccggctcctctctctcttgcaaattctgtagccaaacccatcaaagctcttcctacaccatgac<br>ctctatgctctggggcgacttctatatcttcaacggtcaaccttctattccaacctgaataagaaacgactacaaaaccagccaaatcaccgtcatcaccgtatgctacgaaagttcttgaatctggg<br>tcaccatcttctccgtcgtcgctttcgtcatcagattcatcatctggaaagactttagttaatggaggatcgactgggacttctcttaaggtaaatccatcaccggtggcggttactctaaagacggtgt<br>cggtggtgaatgaaccgtctaaagcctcaattgcttcagcatcaccagggactgatgttctgtatctgtaggctgtatcatctaaggtagtacccattttataattattataaaaactttcatatagagct<br>ccgtttctattatgaatttcatttataaagtttatgtacaaatatcataaaaaaagagaatctttttcatttcttagcatttttgacgaaatttgctattttgttagagtcttttacaccatttgtctccacacctcc<br>gcttacatcaacaccaataacgccatttaatctaagcgcatcaccaacattttctggcgtcagtccaccagctaacataaaatgtaagctttcggggctctcttgccttccaacccagtcagaaatcg<br>agttccaatccaaaagttcacctgtcccacctgcttctgaatcaaacaagggaataaacgaatgaggtttctgtgaagctgcactgagtagtatgttgcagtcttttggaaatacgagtcttttaataa<br>ctggcaaaccgaggaactcttggtattcttgccacgactcatctccatgcagttggacgatatcaatgccgtaatcattgaccagagccaaaacatcctccttaggttgattacgaaacacgccaa<br>ccaagtatttcggagtgcctgaactatttttatatgcttttacaagacttgaaattttccttgcaataaccgggtcaattgttctctttctattgggcacacatataatacccagcaagtcagcatcggaat<br>ctagagcacattctgcggcctctgtgctctgcaagccgcaaactttcaccaatggaccagaactacctgtgaaattaataacagacattttataattattataaaaactttcatataagagatataaaat<br>ttaatatggaagaaataagacaagaaagatacaaccaaatgaaagaagctctaaatagtgttgaaggttataaaggaaaaattgtagcctcagactcagattggtgtttcaaagatcctcaaggca<br>atagaataacagattttgatagtactttaagccagccccgacacccgccaacacccgctgacgcgccctgacgggcttgtctgctcccggcatccgcttacagacaagctgtgaccgtctccgg<br>gagctgcatgtgtcagaggttttcaccgtcatcaccgaaacgcgcgagacgaaaggcaaaaaaaaggctccaaaaggagcctttaattgtatcggtttaccaatgcttaatcagtgaagcccctta<br>tttcagcaatctgtctatttcgttcgtccatagttgcctgactacccgtcgtgtagataactacgatacgggaaggcttaccatctggtccaagtgcggcaatgataccccgtgaaccacgctctcct<br>gccccagatttatcagcaataaaccagccagcaggaagtgccgaacgcagtagtggtcctgcaactttatccgcttcaagccagtctattaattgttgccgtgaagccagagtaagtagttcgcc<br>agttaatagtttgcgtaatgttgttgccattgccgcaggcattgtggtgtctcgctcgtcgtttggtatggcttcattcagttctggttcccaacggtcaaggcgagttacatggtcacccatattgtgca<br>aaaaagcggttaattccttcggtccaccgatggttgtcagaagtaagttcgccgcagtgttatcactcatcgttatggcggcactacataattctcttaccgtcattccgtccgtaaggtgcttttctgt<br>cactggggagtattcaaccaagtcattctgcgaatagtgtattcgtcgtccgagttgctcttgccccgcatctacacgggataatactgctccacatagcagaactttgaaagtgctcatcattggaa<br>aacgctcttcagggcgaaaactctcaaggattttaccgctattcaagtccagttctatgtatcccactcgtgcccccaattggtcctctgcgtcttttactttcacgagcgtttctgggtgagcaaaaac<br>aggaaggcaaaatgccgcaaaaaagggaataagggcgacacggaaatgttgaatactcatactcttcctttttcaatattattgaagcatttatcagggttattgtctgatgagcggatacatatttg<br>aatgtatttagaaaaataaacaaataggggaattaaaaaaaagcccgctcattaggcgggctactagtcggatatagttcctcctttcagcaaaaaacccctcaagacccgtttagaggccccaa<br>ggggttatgctagttattgctcagcggtggcagcagccaactcagcttcctttcgggctttgttagcagccggatcttctagaagaccccgtagaaaagatcaaaggatcttcttgagatcctttttt<br>ctgcgcgtaatctgctgcttgcaaacaaaaaaaccaccgctaccagcggtggtttgtttgccggatcaagagctaccaactctttttccgaaggtaactggcttcagcagagcgcagataccaaat<br>actgttcttctagtgtagccgtagttaggccaccacttcaagaactctgtagcaccgcctacatacctcgctctgctaatcctgttaccagtggctgctgccagtggcgataagtcgtgtcttaccgg<br>gttggactcaagacgatagttaccggataaggcgcagcggtcgggctgaacggggggttcgtgcacacagcccagcttggagcgaacgacctacaccgaactgagatacctacagcgtga<br>gctatgagaaagcgccacgcttcccgaagggagaaaggcggacaggtatccggtaagcggcagggtcggaacaggagagcgcacgagggagcttccagggggaaacgcctggtatcttt<br>atagtcctgtcgggtttcgccacctctgacttgagcgtcgatttttgtgatgctcgtcaggggggcggagcctatggaaaaacgccagcaacgcggcctttttacggttcctggccttttgctggcc<br>ttttgctcacatgttctttcctgcgttatcccctgattctgtggataaccgtattaccgcctttgagtgagctgataccgctcgccgcagccgaacgaccgagcgcagcgagtcagtgagcgagga<br>agcggaagagcgcccaatacgcaaaccgcctctccccgcgcgttggccgattcattaatgcagctggcacgacaggtttcccgactggaaagcgggcagtgagcgcaacgcaatagtacta<br>taatatatgaattacattattaatttaaaaactcgtgatgacctatacataggaagatctatagaaacaaaaagattaataactttcaaatatcagaaaaatatagaaacatgtgataagctcatagacat<br>ataaaaatatgagattcccatctatcttcaccgctgttttgttcgctgcttcttctgctttggctgctccagctaacaccaccaccgaagacgaaaccgctcaaatcccagctgaagctgttatcgact<br>actctgacttggaaggtgacttcgacgctgctgctttgccattgtctaactctaccaacaacggtttgtcttctaccaacaccaccatcgcttctatcgctgctaaggaagaaggtgttcaattggaca<br>agagagaagctgacgcagagaccgaaagtgaaacgtgatttcatgcgtcattttgaacattttgtaaatcttatttaataatgtgtgcggcaattcacatttaatttatgaatgttttcttaacatcgcgg<br>caactcaagaaacggcaggttcggatcttagctactagagaaagaggagaaatactagatgcgtaaaggcgaagagctgttcactggtgtcgtccctattctggtggaactggatggtgatgtc<br>aacggtcataagttttccgtgcgtggcgagggtgaaggtgacgcaactaatggtaaactgacgctgaagttcatctgtactactggtaaactgccggttccttggccgactctggtaacgacgctg<br>acttatggtgttcagtgctttgctcgttatccggaccatatgaagcagcatgacttcttcaagtccgccatgccggaaggctatgtgcaggaacgcacgatttcctttaaggatgacggcacgtaca<br>aaacgcgtgcggaagtgaaatttgaaggcgataccctggtaaaccgcattgagctgaaaggcattgactttaaagaggacggcaatatcctgggccataagctggaatacaattttaacagcca<br>caatgtttacatcaccgccgataaacaaaaaaatggcattaaagcgaattttaaaattcgccacaacgtggaggatggcagcgtgcagctggctgatcactaccagcaaaacactccaatcggt<br>gatggtcctgttctgctgccagacaatcactatctgagcacgcaaagcgttctgtctaaagatccgaacgagaaacgcgatcatatggttctgctggagttcgtaaccgcagcgggcatcacgca<br>tggtatggatgaactgtacaaatgaccaggcatcaaataaaacgaaaggctcagtcgaaagactgggcctttcgttttatctgttgtttgtcggtgaacgctctctactagagtcacactggctcacc<br>ttcgggtgggcctttctgcgtttataggtctcggttccaaggacaatagctcgacgattgaaggtagatacccatacgacgttccagactacgctctgcaggctagtggtggtggtggttctggtg<br>gtggtggttctggtggtggtggttctgctagcgacgtccaggaactgacaactatatgcgagcaaatcccctcaccaactttagaatcgacgccgtactctttgtcaacgactactattttggccaa<br>cgggaaggcaatgcaaggagtttttgaatattacaaatcagtaacgtttgtcagtaattgcggttctcaccccctcaacaactagcaaaggcagccccataaacacacagtatgtttttgaatcctaa<br>taatggccgcaaaaaaaaaaaaaaaaaaaaaaaaaaaaaaaaaaaaaaaaaaaaaaaaaaaaaaaaaaaaaaaaacctgtcaccggatgtgttttccggtctgatgagtccg<br>tgaggacgaaacaggactag |
